# The drought-responsive *ZmFDL1* gene regulates cuticle biosynthesis and cuticle-dependent leaf permeability

**DOI:** 10.1101/2020.03.30.014332

**Authors:** Giulia Castorina, Frédéric Domergue, Matteo Chiara, Massimo Zilio, Martina Persico, Valentina Ricciardi, David Stephen Horner, Gabriella Consonni

## Abstract

In higher plants, the outer surface of the aerial parts is covered by the cuticle, a complex lipid layer that constitutes a barrier against damages caused by environmental factors and provides protection against non-stomatal water loss. We show in this study that cuticle deposition, during the juvenile phase of in maize (*Zea mays*) plant development, and cuticle-dependent leaf permeability are controlled by the MYB transcription factor *ZmMYB94*/*FUSED LEAVES1* (*ZmFDL1*).

Biochemical analysis showed that in *fdl1-1* mutant seedlings at the coleoptile stage both cutin and wax biosynthesis and deposition were altered. Among cutin compounds, ω-hydroxy fatty acids and polyhydroxy-fatty acids were specifically affected, while the reduction of epicuticular waxes, was mainly observed in primary long chain alcohols, and to a minor extent, long-chain wax esters.

Transcriptome analysis allowed the identification of novel candidate genes involved in lipid metabolism and the assembly of a proposed pathway for cuticle biosynthesis in maize. Lack of ZmFDL1 affects the expression of genes located in different modules of the pathway and correspondence between gene transcriptional variations and biochemical defects have been highlighted.

A decrease in cuticle-dependent leaf permeability was observed in maize seedlings exposed to drought as well as ABA treatment, which implies coordinated changes in the transcript levels of *ZmFDL1* and associated genes. Overall, our results suggest that the response to water stress implies the activation of wax biosynthesis and the involvement of by both ZmFDL1 and ABA regulatory pathways.

**One-Sentence Summary:** Cuticle biosynthesis and cuticle-mediated drought-response during the juvenile phase of maize plant growth, are regulated by the MYB transcription factor fused leaves1 (ZmFDL1) and influenced by ABA.

## INTRODUCTION

In higher plants, the outer surface of aerial parts, including vegetative organs, flowers, fruits, seeds and pollen grains, is constituted by a continuous hydrophobic layer termed cuticle, which consists of two major components, the polymer cutin and cuticular waxes. Cutin is a polymer of C16 to C18 hydroxylated fatty acids, that are directly cross-esterified to each other or via glycerol backbone (Fich et al., 2016). Cutin monomers are synthetized in the endoplasmic reticulum (ER), through different reactions, including the esterification of fatty acids to coenzyme A (CoA), ω-hydroxylation and further oxidation, glycerol-3-phosphate transacylation, and export through the cell wall to the surface where polymerization occurs (Fich et al., 2016). Cuticular waxes are constituted by a complex mixture of very-long-chain fatty acids (VLCFAs), with more than 20 carbon atoms, and their derivatives, which include alcohols, aldehydes, alkanes, ketones and wax esters. They also include variable amounts of cyclic compounds, such as triterpenoids and phenylpropanoids (Bernard and Joubès, 2013; Lee and Suh, 2015a). The first phase of wax biosynthesis consists in the elongation of C16 and C18 fatty acids produced in the plastids to very-long-chain fatty acids (VLCFAs), with a chain length between C22 and C38, by elongase (FAE) complexes in the endoplasmic reticulum (Haslam and Kunst, 2013). Wax components are then produced through two different pathways. Primary alcohols and esters are produced by the alcohol forming pathway, also termed acyl-reduction pathway (Rowland et al., 2006; Li et al., 2008), while alkanes, secondary *n-*alcohols and ketones are produced by the alkane forming pathway, also called decarbonylation pathway (Bernard et al., 2012). All wax compounds are synthesized in the epidermal cell layer (L1) and are secreted to cover the cell wall of epidermal cells, where they are embedded in the cutin and deposited on the cuticle surface as films or wax crystals. Waxes forming the outer layer of plant tissues are called epicuticular waxes.

Cuticle formation prevents post genital fusions among organs that grow very tightly appressed to each other when enclosed in vegetative shoots or within buds (Ingram and Nawrath, 2017). In addition, cuticle constitutes a constant barrier against damages caused by environmental abiotic and biotic factors, including UV light, temperature changes, pest and pathogens and a primary waterproof barrier that controls non-stomatal water loss and gas exchange from leaves (Yeats and Rose, 2013). Several studies in various plants have shown that changes in the amount and/or distribution of cuticular waxes lead to alterations in cuticular permeability. In tomato, it was shown through mechanical and genetic manipulations of the cuticular components that aliphatic constituents of the intracuticular wax layer have a key role in limiting transpiration rate across epidermis (Vogg et al., 2004). In *Sorghum bicolor*, an irradiate bloomless mutant with a high reduction in epicuticular waxes appeared to have a higher rate of epidermal permeability and night-time water loss (Burow et al., 2008). In rice, impairments in the organization of crystal waxes on the leaf surfaces of the *wilted dwarf and lethal 1* (*wdl1*) mutant were correlated with a 2.3-fold increase in the transpiration rates and higher rates of water loss (Park et al., 2010). A positive correlation between an increase of wax amount and tolerance to drought stress was reported in Arabidopsis (*Arabidopsis thaliana*) (Aharoni et al., 2004; Kosma et al., 2009; Cui et al., 2016). Similarly, alkanes and primary alcohols were increased by drought treatment in two Australian wheat cultivars. Their study allowed the characterization of two MYB transcription factors that were activated by drought and in turn, stimulated the expression of cuticle biosynthesis-related genes (Bi et al., 2016). In rice, the *OsGL1-6* gene, highly similar to Arabidopsis *AtCER1* and involved in cuticular wax accumulation, was shown to be activated by drought (Zhou et al., 2015). Finally, in maize seedlings, both cuticle permeability and drought sensitivity were shown to be increased by a mutation in the *GLOSSY6* (Zm*GL6*) gene, the product of which is putatively involved in intracellular trafficking of cuticular waxes (Li et al., 2019).

In Arabidopsis, deposition of cuticular waxes in both leaves and stems is regulated by the AtMYB96, AtMYB30 and AtMYB94 transcription factors, that collectively modulate the expression of wax biosynthetic enzymes (Raffaele et al., 2008; Seo et al., 2011; Lee and Suh, 2015a). ABA, drought, and high salinity activate the expression of *AtMYB96* (Seo et al., 2009) which in turn mediates the activation of cuticle biosynthetic genes to increase drought tolerance (Seo et al., 2011). Similarly, the expression level of *AtMYB94* is also increased by drought, and transgenic Arabidopsis lines overexpressing *AtMYB94* showed an increased accumulation of cuticular waxes and, under drought conditions, a reduction in the rate of cuticular transpiration in leaves (Lee and Suh, 2015b). We showed in a previous work that the maize *ZmFDL1*/*MYB94* transcription factor is closely related to AtMYB96, AtMYB30 and AtMYB94 and is responsible for cuticle mediated post-genital organ separation during embryo development and early phases of seedling growth. Lack of *ZmFDL1* activity, in the *fused leaves1-1* (*fdl1-1*) recessive mutant, specifically affects seedling development at early developmental stages and results in organ fusions, due to the lack of cuticular material in the boundary between organs, and irregular distribution of wax crystals on young leaf epidermal surface (La Rocca et al., 2015).

To further gain insights into the role of the *ZmFDL1/MYB94*, we compared in this study the cuticle composition of mutant and wild type seedlings and analysed the impact of the mutation on the transcriptome during early phases of seedling development. We also investigated *ZmFDL1/MYB94* involvement in controlling cuticular permeability and in mediating drought stress response in maize seedlings. The isolation of genes involved in biosynthesis and transport of cuticular waxes and responsive to drought are of particular interest for crop breeding. The detailed functional characterization of the maize *ZmFDL1*/*MYB94* regulatory gene proposed in this work will contribute to unravel the genetic-molecular mechanisms at the basis of the cuticle-mediated drought stress tolerance in an important cereal crop such as maize.

## RESULTS

### ZmFDL1 regulates cuticle deposition in a phase-dependent manner

The most evident defects observed in *fdl1-1* homozygous mutants were the irregular coleoptile opening and presence of fusions between coleoptile and first leaf (Fig. 1A; La Rocca et al., 2015). To characterize the effects of the *fdl1-1* mutation on cuticle composition, cutin and epicuticular waxes were extracted from mutant and wild type seedlings and analysed by gas chromatography (Domergue et al., 2010; Bourdenx et al., 2011).

**Figure 1.**
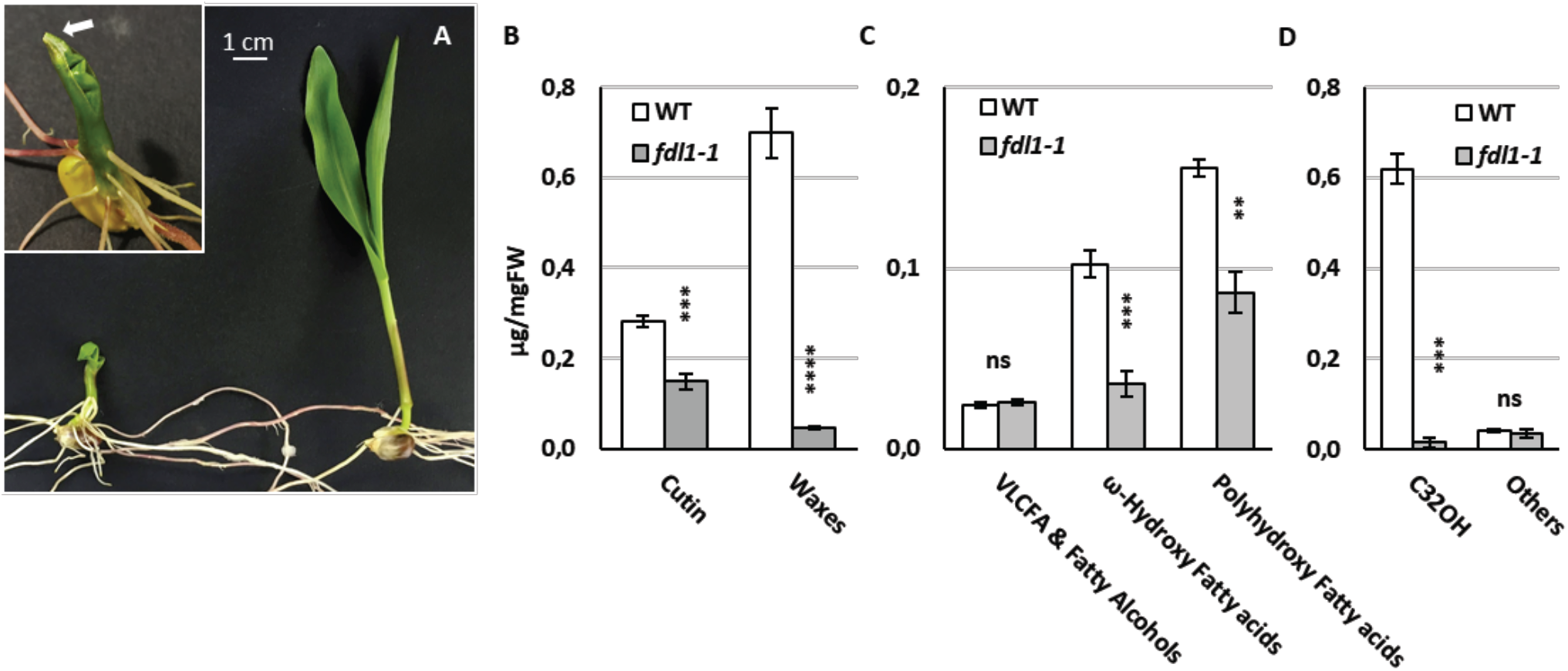
Cuticle related defects in *fdl1-1* homozygous mutant. (A) Representative phenotype of wild type (WT, right) and 10-days-old *fdl1-1* mutant (left) seedlings. Inset: the white arrow indicates the fusion between the coleoptile and the first leaf of a *fdl1-1* plant. Total cutin and wax loads (B), Cutin composition (C), and Wax composition (D) in *fdl1-1* mutant and wild type (WT) seedlings at the coleoptile developmental stage. The data represent the means ± SE of five biological replicates. Significant differences were assessed by Student’s t-test (* = P < 0.05, ** = P < 0.01, *** = P < 0.001 and **** = P < 0.0001).

At the coleoptile stage, the total cutin and wax loads of *fdl1-1* mutants were considerably reduced compared with those of wild type seedlings, but not completely abolished (Fig. 1B). In *fdl1-1* coleoptiles, the reduction in total waxes was more severe than that observed for cutin (94% and 47% decreases, respectively; Fig. 1B). These differences progressively decreased during seedling development and at the third leaf stage, total amounts of cutin (Supplemental Fig. S1A) and waxes (Supplemental Fig. S1B, S1C, S2M-T) were similar in *fdl1-1* and wild-type plants. Reduction in cutin content was mainly due to decreases in ω-hydroxy fatty acids and polyhydroxy-fatty acids (Fig. 1C and Supplemental Fig. S2E-L). The ω-hydroxy fatty acids content was more impaired than the polyhydroxy-fatty acids content (65% and 44% decreases, respectively) in the coleoptiles of the homozygous *fdl1-1* mutant (Fig 1C). Such differences were detected up to the second leaf stage of the seedling development (Supplemental Fig. S2E, F, I and J). Loss of ZmFDL1 activity had no substantial effect on the very-long-chain fatty acids (VLCFA) and fatty alcohols (Fig 1C) as only a few monomers were significantly different from wild type at the early stage of seedling development (Supplemental Fig. S2A, B).

The decrease in epicuticular wax load (Fig. 1B and Supplemental Fig. S1, S2) was mainly due to reduction in primary long-chain alcohols (Fig. 1D and Supplemental Fig. S1, S2), which represent the major components of maize seedling waxes and are dominated by the C32 primary alcohol isomer. In coleoptiles, the C32 primary alcohol content was decreased by 98% in *fdl1-1* when compared to wild type (Fig. 1D). The amounts of other minor primary long-chain alcohols with 30 or more carbons (C31 to C34OH) were also lower in *fdl1-1*, whereas the amounts of fatty alcohols with less than 30 carbons (C28 and C26OH) showed the opposite trend and were increased in *fdl1-1* with respect to wild type seedlings (Supplemental Fig. S2Q, R). These differences disappeared with seedling development, and in the third leaf, the amounts of primary alcohol were similar in *fdl1-1* mutant and wild-type seedlings (Fig S1C; S2 Q-T). Alkanes and aldehydes were also analysed (Supplemental Fig. S2M-P) and found to be reduced only in the early stages of seedling development (Supplemental Fig. S2M, N). In particular, the C32 n-aldehyde was nearly absent in *fdl1-1* coleoptiles whereas alkanes were less affected (Supplemental Fig. S2M). Finally, minor long-chain wax esters (WE) were detected at the coleoptile stage and their content was much lower in the homozygous *fdl1-1* mutant than in wild type seedlings (Supplemental Fig. S1D). Altogether these data suggested that ZmMYB94 is a regulatory component of both cutin and wax biosynthesis and deposition. Moreover, as previously observed for visual mutant traits, ZmMYB94 related biochemical defects undergo a progressive reversion during seedling development, further confirming that ZmMYB94 activity occurs in a specific developmental phase (La Rocca et al., 2015).

### Leaf permeability is altered in the *fdl1-1* mutant

Cuticle-dependent leaf permeability was assessed by means of the chlorophyll leaching assay. The second leaf of *fdl1-1* homozygous seedlings displayed a higher permeability compared to non-mutant B73 siblings (Fig. 2A). Differences observed between the mutant and wild type leaf permeability progressively decreased in the second, third and fourth leaves (Fig. 2B). The differences were maintained in different genetic backgrounds, as observed in F_2_ progenies obtained after introgressing the mutation in the H99 and Mo17 inbred lines (Fig. 2C). To further understand the impact of the increased leaf epidermal permeability, thermography images of second fully expanded leaves of wild type (Fig. 2E) and *fdl1-1* homozygous mutant (Fig. 2F) seedlings were analysed. The *fdl1-1* presented a reduced leaf temperature of about 1.5°C compared to its control (Fig. 2D). Moreover, a water loss time course experiment was performed on 10-old-days seedlings, by estimating the loss of weight with respect to the initial seedling fresh weight. The resulting profiles showed that homozygous *fdl1-1* had a higher water loss rate compared to the wild type plants (Fig. 2G). We excluded that the observed differences in epidermal leaf permeability, leaf temperature and water loss values could be due to alterations in the stomatal index (Supplemental Fig. S4A, B).

**Figure 2.**
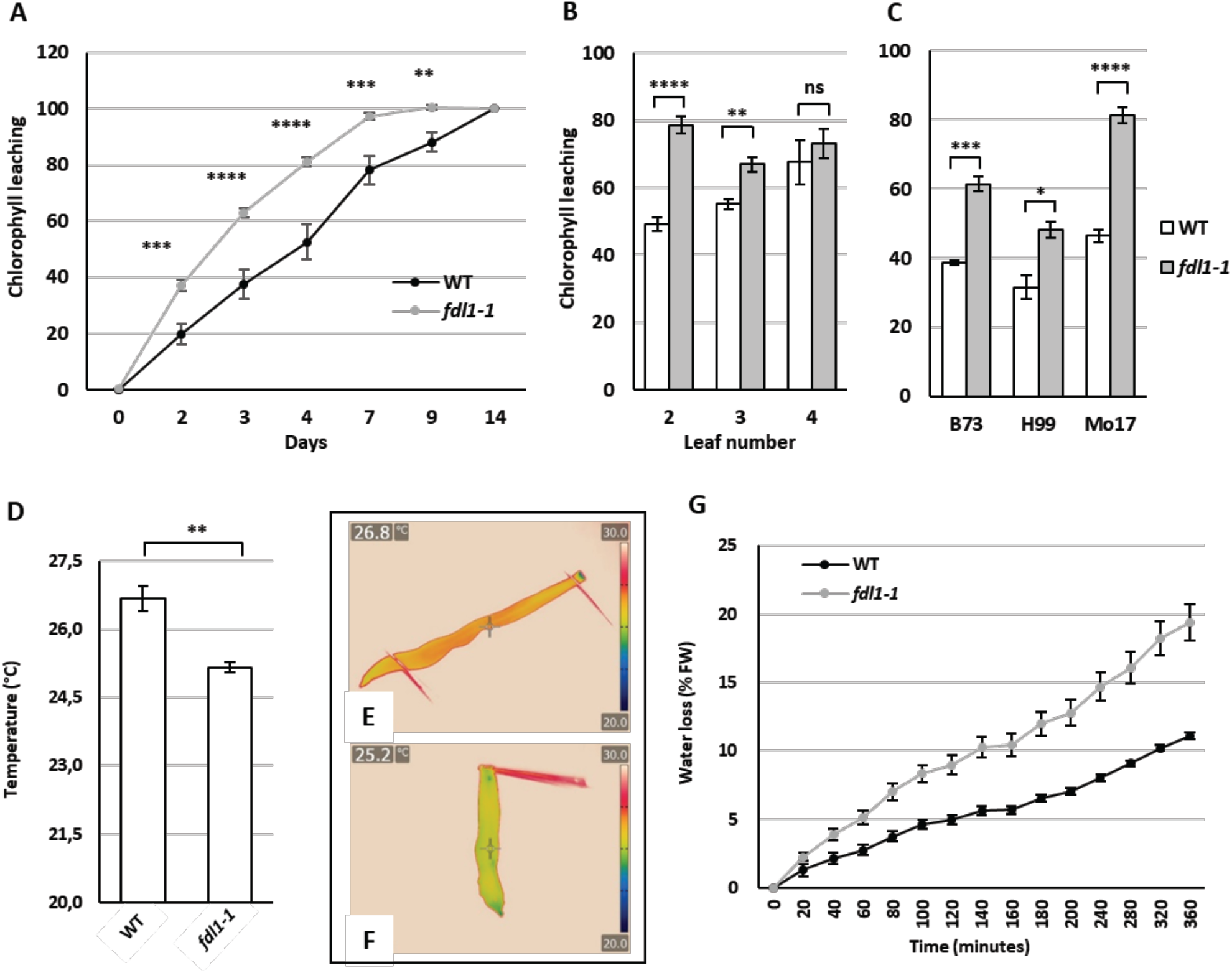
Cuticle-dependent leaf permeability in the *fdl1-1* mutant. The chlorophyll leaching assay was performed on (A) the second fully expanded leaf of 14-days-old *fdl1-1* and B73 wild type (WT) control plants; (B) in the second, third and fourth fully expanded leaf of *fdl1-1* mutant compared to the wild type (WT) control plants and (C) in the second fully expanded leaf of *fdl1-1* mutant and wild type (WT) plants in different genetic background. The (B) and (C) panels represent the date at day 3 of the chlorophyll leaching assay. Values represent the mean ± SE of a minimum of five (A-C) replicates. The comparison was made at each time point between wild type and homozygous *fdl1-1* genotypes. (D) The temperature of the second fully expanded leaf in 14-old-days wild type (WT) and *fdl1-1* homozygous plants. Values represent the mean ± SE of five biological replicates. Representative images of the leaf temperature acquired with the thermographic camera in (E) wild type and (F) homozygous *fdl1-1* plants. (G) Percentage of water loss in detached 10-old-days homozygous *fdl1-1* and wild type (WT) seedlings. Values represent the mean ± SE of 13 and 21 biological replicates for wild type (WT) and *fdl1-1* homozygous plants, respectively. Significant differences were assessed by Student’s t-test (* = P < 0.05, ** = P < 0.01, *** = P < 0.001, **** = P < 0.0001, ns = not significant).

### Transcriptome profile of the *fdl1-1* mutant

To investigate the regulatory network associated with ZmFDL1, cDNA libraries were produced from coleoptiles of *fdl1-1* and wild type seedlings and sequenced on HiSeq™ 2000 platform. Six biological replicates, three for the *fdl1-1* mutant and three for the wild type background respectively, were analysed. Statistics concerning the total number of reads produced and the proportion of reads assigned to gene models according to the Zm00001d annotation of the ZmB73_RefGen_v4 reference assembly (Jiao et al., 2017) are reported in Supplementary Table S1. Identification of differentially expressed genes (DEGs) was performed by the latest versions of DESeq2 (Love et al., 2014) and Limma (Ritchie et al., 2015). A total of 2213 and 2399 genes were considered differentially expressed by Limma and DESeq2 respectively when a significance cutoff FDR of 0.05 was applied. Importantly, 1639 genes were differentially expressed according to both methods and were considered for subsequent analyses. Of these 612 were down-regulated and 1027 were up-regulated (Fig. 3A).

**Figure 3.**
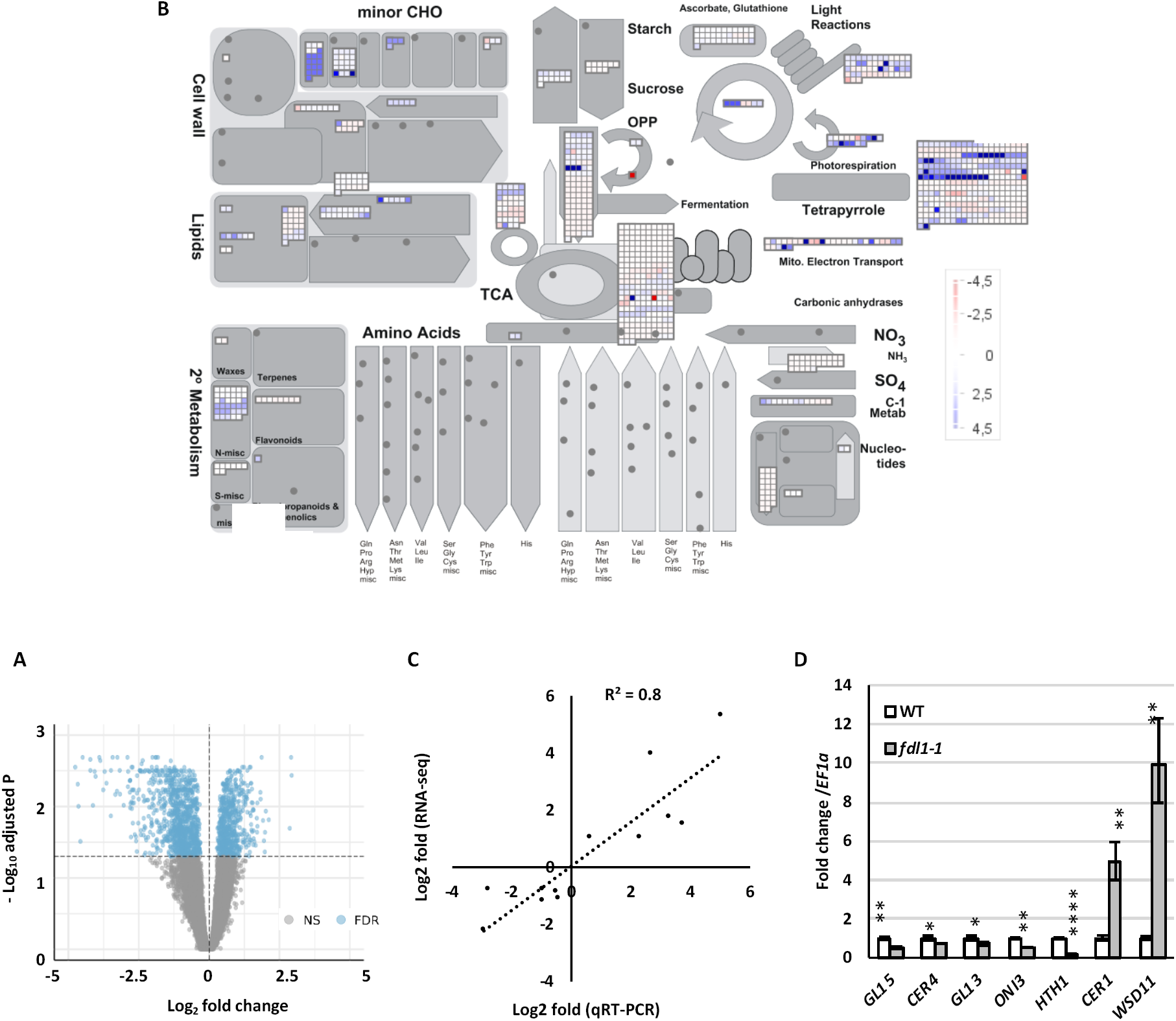
Functional enrichment of metabolic pathways and cuticle related genes. (A) Volcano plot of differentially expressed genes (DEGs) in *fdl1-1* compared to wild type control plants. Blue dots denote up-regulated and down-regulated DEGs. Grey dots represent genes with no significant P value. (B) Functional enrichment of metabolic pathways. The DEGs have been assigned to different cellular components through the Gene Ontology analysis. MAPMAN was used to have a general overview of differentially expressed transcripts involved in different metabolic pathways and cellular processes. ZmFDL1-mediated changes in different metabolic processes in seedlings are illustrated (blue, induced; red, repressed). (C) Quantitative Real-Time PCR (qRT-PCR) validation of differentially expressed genes (DEGs) characterized by RNA-seq. Correlation of Log_2_ fold change data obtained using Real-Time PCR (y axis) and with RNA-Seq analysis (x axis). (D) Gene expression level of putative cuticle related genes, analysed by qRT-PCR, in *fdl1-1* and wild type (WT) control seedlings at the coleoptile developmental stage. Values represent the mean fold change variations ± SD of four biological replicates. Comparison has been made between wild type and homozygous fdl1-1 genotypes. Significant differences were assessed by Student’s t-test (* = P < 0.05, ** = P < 0.01, *** = P < 0.001, **** = P < 0.0001, ns = not significant).

Functional enrichment analyses of differentially expressed genes, according to the Gene Ontology (GO) annotation of maize gene models reported by (Wimalanathan et al., 2018), was performed in order to identify biological processes and pathways that are differentially modulated in the *fdl1-1* mutant. As reported in the Supplementary Tables S4 and S5 more than 200 ontology terms were significantly enriched in both up and down-regulated genes, after FDR correction, suggesting that the depletion of ZmFDL1 is likely to elicit a complex regulatory response. Importantly, ontology terms related to phenotypic traits observed in the *fdl1-1* mutant, including: “GO:0010025: wax biosynthetic process”, “GO:0005618: cell-wall”, “GO:0009414: response to water deprivation” and “GO:0019216: regulation of lipid metabolic process” are highly enriched in both sets of up and down-regulated genes (Fig. S3B, C), suggesting that differentially expressed genes identified by our analyses are likely to be involved in the modulation of wax/cutin biosynthesis and deposition in maize. The most significantly enriched GO terms associated with up-regulated genes include “GO:0042542: response to hydrogen peroxide”, “GO:0009408: response to heat”, “GO:0010286: heat acclimation”, “GO:0009651: response to salt stress” and similar (Supplementary Table S3), suggesting that the majority of the genes up-regulated in the *fdl1-1* mutant, might be involved in the response to osmotic and metabolic stress caused by impaired *fdl1* activity. On the other hand, down-regulated genes are highly enriched in ontology terms associated with chloroplast and pigments, including for example: “GO:0009570: chloroplast stroma”,” GO:0009941: chloroplast envelope”, “GO:0016117: carotenoid biosynthetic process” and “GO:0015995: chlorophyll biosynthetic process”.

Consistent with the enrichment pathways recovered from Gene Ontology analyses, MapMan (Schwacke et al., 2019) (Fig. 3B) ontology analyses of metabolic pathways in the *fdl1-1* mutant, suggested a high enrichment of metabolic terms related to organelles, including for example: “.RNA processing.organelle machinery”, “.RNA biosynthesis.organelle machinery”, “.Protein biosynthesis and “.organelle machinery”, pigments: “.External stimuli response.light”, “.External stimuli response.light.UV-A/blue light” and “blue-light mediated photoperception”, but also of terms related to cell wall organization and lipid biosynthesis such as: “.Cell wall organisation.cutin and suberine” and “.Lipid metabolism”.

Functional enrichment analyses of KEGG pathways (Moriya et al., 2007; Okuda et al., 2008) identified the “zma00195: Photosynthesis” and “zma04712: Circadian rhythm” pathways as those with the most statistically significant enrichment in up- and down-regulated genes, respectively. Fatty acid biosynthesis occurs in the plastid (Fig. 4) and conceivably an impairment of the downstream usage of the early cutin and wax monomer precursors could lead to alterations in the chloroplast physiology and produce an altered photosynthetic activity.

**Figure 4.**
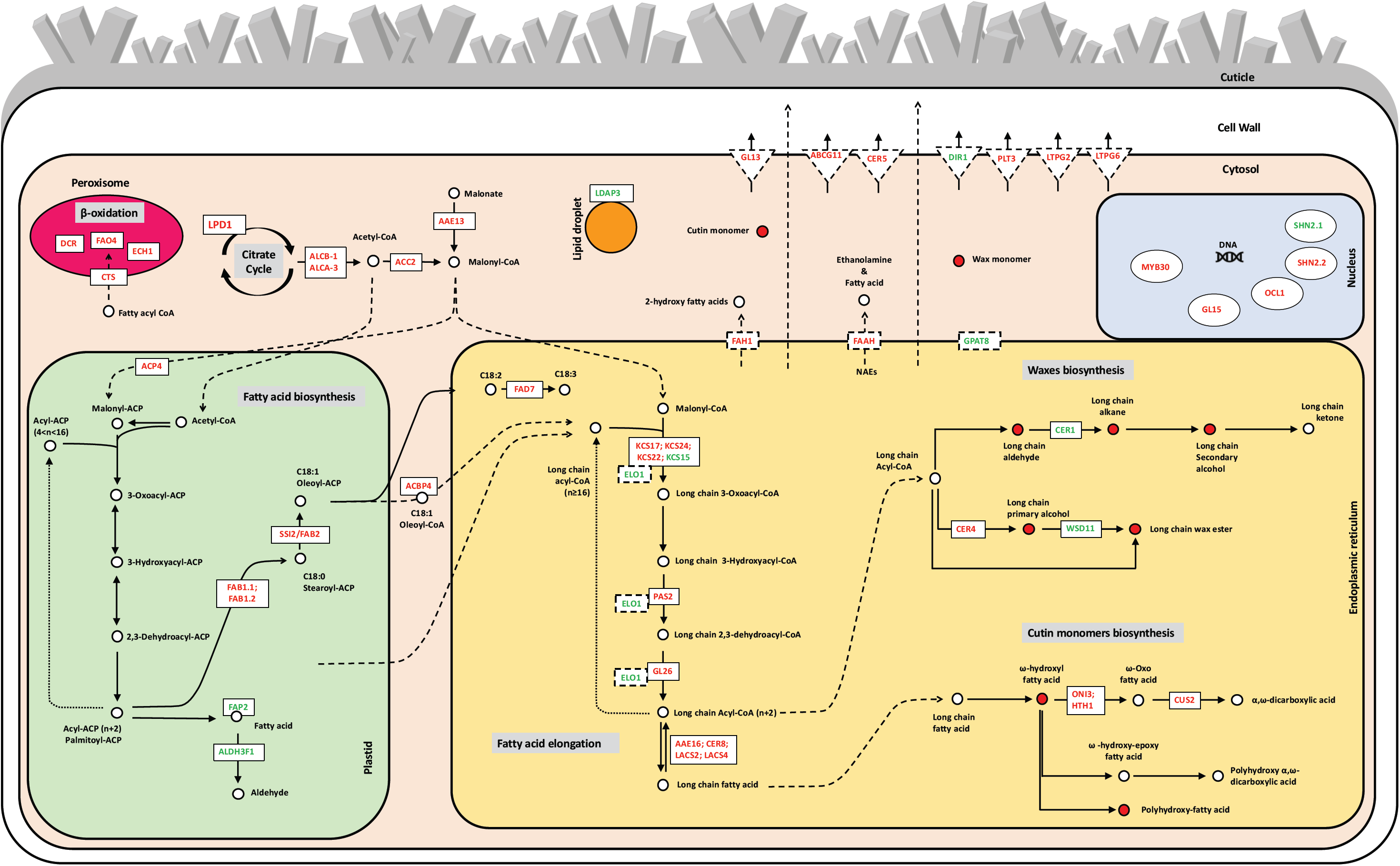
Pathways involved in cuticle biosynthesis. Schemes were designed using pathways available at KEGG database. Transcript regulation, Citrate cycle and Acetyl-CoA biosynthesis, fatty acid biosynthesis, fatty acid elongation, wax biosynthesis, cutin biosynthesis and cuticle monomers transport are represented in the corresponding cell compartments. Solid black arrows indicate enzymatic reaction steps and gene symbols are referred to DEGs in specific steps. Intermediates are shown as circles that are red coloured when their amount is lower in *fdl1-1* mutant compared to the WT plants. Dashed black arrows indicate the movement of specific compounds into or towards a cell compartment. Down and up-regulated genes (Supplemental Table S1), are represented with red and green coloured symbols, respectively. Genes encoding soluble enzymes are represented within rectangles with a continuous line, whereas genes encoding transmembrane enzymes are represented with rectangles with a dashed line. Transporters of cuticle monomers are represented with triangles.

More interestingly and in line with previous observations, the pathways related to cutin and fatty acid metabolism among were the most enriched pathways in both genes significantly down-regulated (zma01212: Fatty acid metabolism, zma00061: Fatty acid biosynthesis and zma00073: Cutin, suberine and wax biosynthesis), and up-regulated (zma00561: Glycerolipid metabolism). All in all, the high level of agreement observed between the results of our functional enrichment analyses, which were performed according to three independent ontologies, are highly consistent with the idea that genes differentially modulated in the *fdl1-1* mutant are likely to mediate the phenotypic traits observed in this developmental stage, as demonstrated by the systematic enrichment of ontology terms associated with wax, cutin and lipid metabolism in general.

Accordingly, we performed a careful manual annotation of our list of differentially expressed genes to identify candidate genes directly involved in cutin metabolism. Based on a combination of Gene Ontology, KEGG pathway and orthology predictions with *Arabidopsis thaliana* and *Oryza sativa*, as available from the phytozome annotation of the ZmB73_RefGen_v4 gene models, and the ARALIP Website (Li-Beisson et al., 2013) we identified 79 candidate DEGs which were tentatively assigned to 9 distinct lipids and cuticle-related biosynthetic processes, defined by expert manual curation (Supplemental Table S1). These genes were mainly related to the biosynthesis and the transport of secondary metabolites but also to the regulation of the gene expression. According to our careful manual annotation, 15 DEGs were assigned generically to the lipid metabolism, 9 DEGs to the glycerolipid metabolism and 5 DEGs to the glycerophospholipid metabolism. Remaining genes were assigned to the putative biosynthetic pathways described below.

The Zm00001d009212 (*ZmLPD1*); Zm00001d040603 (*ZmACLB-1*); Zm00001d048627 (*ZmACLA-3*); Zm00001d004125 (*ZmACC2*) and Zm00001d043376 (ZmAAE13) genes were associated with citrate cycle, Acetyl-CoA and Malonyl-CoA biosynthesis, respectively. The Zm00001d041701 (*ZmACP4*), Zm00001d004019 (*ZmSSI2*/*ZmFAB2*), Zm00001d006866 (*ZmFAB1.1*), Zm00001d022144 (*ZmFAB1.2*), Zm00001d017418 (*ZmALDH3F1*) and Zm00001d047743 (*ZmFAD7*) genes were related to biosynthesis and desaturation of fatty acid. The Zm00001d008622 (*ZmGL26*); Zm00001d039856 (*ZmPAS2*); Zm00001d029350 (*ZmKCS24*); Zm00001d037328 (*ZmKCS17*); Zm00001d044579 (*ZmKCS22*/*ZmCER60*); Zm00001d039053 (*ZmKCS15*) and Zm00001d033637 (*ZmELO1*) genes were associated with the fatty acid elongation pathway.

The Zm00001d024723 (*ZmCER8*); Zm00001d053127 (*ZmLACS2*); Zm00001d045295 (*ZmLACS4*); Zm00001d034832 (*ZmAAE16*); Zm00001d017251 (*ZmCER1*); Zm00001d049950 (*ZmMSH1*/*ZmCER4*); Zm00001d043853 (*ZmWSD11*); Zm00001d032284 (*ZmHTH1*); Zm00001d020238 (*ZmONI3*); Zm00001d051923 (*ZmCUS2*); Zm00001d044136 (*ZmGPAT8*); Zm00001d034319 (*ZmFAH1*), Zm00001d004817 (*ZmFAAH*), Zm00001d010855 (*ZmLDAP3*) and Zm00001d031893 (*ZmHCT12*/*ZmDCR*); Zm00001d009182 (*ZmECH1*), Zm00001d051130 (*ZmFAO4*) genes were assigned to the suberine, wax and cutin biosynthesis (in ER) and b-oxidation (in peroxisome) pathway, respectively.

In addition, the Zm00001d013960 (*ZmABCG11*); Zm00001d010426 (*ZmCER5*); Zm00001d011091 (*ZmCTS*); Zm00001d048718 (*ZmACBP4*); Zm00001d039631 (*ZmGL13*); Zm00001d018278 (*ZmCHI4*/*ZmFAP2*); Zm00001d034513 (*ZmLTPG6*); Zm00001d012535 (*ZmLTPG2*); Zm00001d043049 (*ZmPLT3*) and Zm00001d005146 (*ZmDIR1*) genes have a role in binding and transport of the fatty acid, cutin and wax monomers. Finally, the Zm00001d040090 (*ZmOCL1*); Zm00001d020457 (*ZmMYB30*), Zm00001d046621 (*ZmGL15*); Zm00001d035835 (*ZmSHN2.1*); Zm00001d026486 (*ZmSHN2.2*) genes codify for regulatory proteins. The data obtained, provided a good overview and valuable information for investigating the ZmFDL1-dependent cuticle regulation, biosynthesis and transport in maize (Fig. 4 and Supplemental Table S1).

To validate the reproducibility of the gene expression data as obtained by the RNA-Seq analysis, expression patterns of 12 DEGs were investigated by quantitative Real-Time PCR. Notably, RNA-Seq expression data were confirmed for all the DEGs considered and displayed high levels of correlation with Real-Time PCR data (R ^2^ = 0.8) (Fig. 3C). Among the 12 DEGs selected for validation, 7 DEGs, which were differentially expressed in *fdl1-1* compared to wild type seedlings at the coleoptile developmental stage, are putative cuticle-related genes (Fig. 3E).

### Drought affects cuticular permeability and the expression of cuticle-related genes

Wild type B73 seedlings were grown under either drought stress (LW), which was applied by withholding irrigation and maintaining RSWC at 30%, or normal water (NW) conditions. After 14 days, the morphological parameters of seedlings grown under restricted water were similar to those of control plants (Fig. 5A), indicating that the applied stress was mild. We performed chlorophyll leaching assays and observed that treated seedlings (LW) responded to the low water content by modifying the cuticle-dependent leaf permeability, which resulted lower compared to the control plants grown under NW condition (Fig. 5B). The same response was observed in wild type seedlings treated by applying a 1µM ABA solution directly to the roots up to 14 DAS. Chlorophyll leaching assay on the second leaf revealed, also in this case, that epidermal permeability decreased in ABA-treated plants compared to mock-treated control seedlings (Fig. 5C).

**Figure 5.**
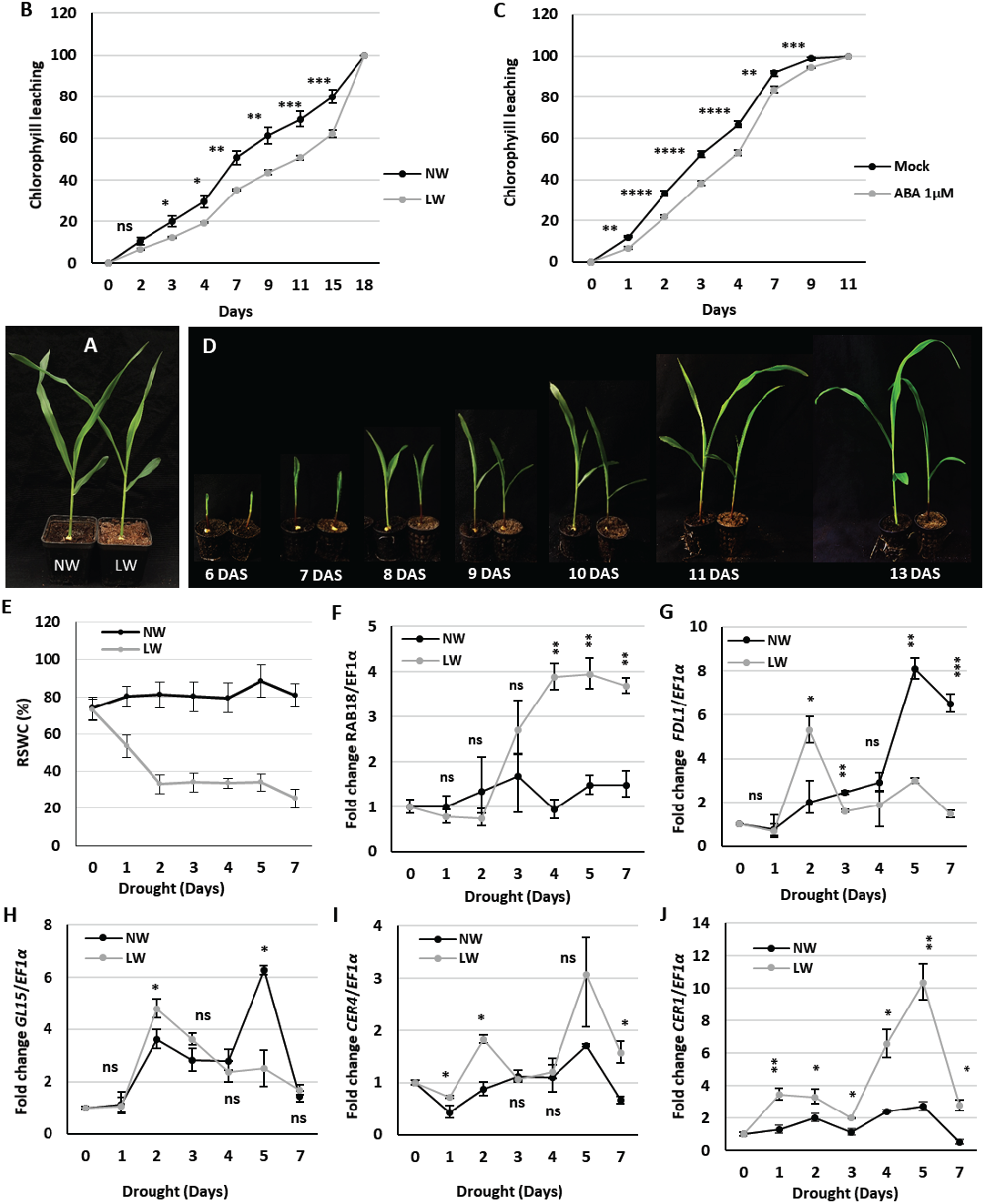
Drought modulates cuticle-dependent leaf permeability and the expression of cuticle related genes. (A) Representative images of 18-days-old B73 wild type seedlings grown under normal (NW) or low water (LW) conditions for 13 days. Leaf permeability of (B) B73 wild type plants grown for 13 days under low-watered (LW) or normal-watered (NW) conditions and (C) B73 wild type plants treated with control (Mock) or 1µM ABA solution. Values represent the mean ± SE of ten (B, C) replicates. Comparison was made at each time point between LW or ABA treated and control plants. (D) Representative images of B73 wild type seedlings grown under NW (left) or LW (right) conditions. Water stress was applied to 6-days-old seedlings and was monitored through the measurement of (E) the Relative Soil Water Content (RSWC) and (F) the expression analysing of the drought stress marker gene *ZmRAB18*. (G-J) Pattern of (G) *ZmFDL1*, (H) *ZmGL15*, (I) *ZmCER4* and (J) *ZmCER1* transcript accumulation analysed by qRT-PCR in the second leaves of wild type plants grown under low water (LW) or normal (NW) conditions. Values represent the mean fold change variations ± SD of three biological replicates. Comparison was made at each time point between LW and control (NW) plants. Significant differences were assessed by Student’s t-test (* = P < 0.05, ** = P < 0.01, *** = P < 0.001, **** = P < 0.0001, ns = not significant).

To investigate transcriptomic changes induced by the drought stress stimuli, maize seedlings were grown under normal-watered (NW) condition until 5 days after germination (DAS) and the next day the drought stress was imposed (Fig. 5D). To apply moderately stronger stress, RSWC was maintained at 30% of the total (Fig. 5E) for a shorter period. As a consequence, after 7 days (13 DAS) treated plants were visibly stunted compared to control seedlings (Fig. 5D). Every day, the second leaf from LW treated and NW control seedlings were sampled and gene expression analysis was performed for every time point.

Expression levels of the drought stress marker *ZmRAB18* (Mao et al., 2015), which resulted higher in seedlings grown under LW compared to those grown under NW conditions (Fig. 5F), confirmed the effectiveness of the drought treatment. The expression profile of *ZmFDL1*, analysed in the second leaf of seedlings grown under NW condition, revealed that *ZmFDL1* transcript accumulates during leaf expansion to reach a peak around 11 DAS (Fig. 5G, black line). In LW condition (Fig. 5G, grey line), *ZmFDL1* transcript profile was different since the peak of maximum expression was detected earlier, after 2 days of drought stress (Fig. 5D, 8DAS), exactly when the RSWC reached the 30% of the total (Fig. 5E). At subsequent time points, the *ZmFDL1* transcript level decreased to be lower in LW than in NW growth conditions, eventually.

The *ZmGL15* transcription factor expression profile nicely mirrored that observed for *ZmFDL1* (Fig. 5H). An increase in transcript accumulation was detected during leaf growth (Fig. 5H, black line). Compared to the control condition, *ZmGL15* was upregulated after 2 days of drought stress and then downregulated, after 5 days of stress (Fig. 5H, grey line). On the other hand, the expression pattern of the *ZmCER4, ZmCER1* and *ZmWSD11* genes showed a trend of upregulation under LW. After 2 and 5 days of stress, *ZmCER4* was significantly upregulated (Fig. 5I, grey line) compared to NW, whereas the transcript level of *ZmCER1*, and *ZmWSD11* which were consistently more expressed in seedlings grown under LW compared to NW condition, reached a peak at 5 and 7 days respectively after the initiation of the stress (Fig. 5J, Supplemental Fig. S4C, grey line).

The expression profiles of *ZmGL13, ZmONI3* and *ZmHTH1* in control conditions decreased throughout leaf development (Supplemental Fig. S4D-F, black line). The *ZmGL13* gene was slightly upregulated after 2 days of drought (Supplemental Fig. S4D, grey line) at the same time point when both *ZmFDL1, ZmGL15, ZmCER4, ZmCER1* and *ZmWSD11* were upregulated (Fig 5G-J, Supplemental Fig. S4C). Instead, the transcripts levels of *ZmONI3* and *ZmHTH1* were less abundant in seedlings grown in LW condition compared to control plants (Supplemental Fig. SE, F, grey line).

### ABA treatment affects *ZmFDL1* and wax-related genes expression

We previously showed that plants responded to ABA treatment by reducing leaf permeability. To investigate the possible involvement of ABA in the control of cuticle-dependent leaf permeability in maize seedling, we analyse a maize mutant in the *VIVIPAROUS1* (*ZmVP1*) gene (*vp1-1*). The transcription factor VP1, is the orthologue of the Arabidopsis *ABA INSENSITIVE3* (*AtABI3*) gene (Giraudat et al., 1992), and mutants in this gene have been reported to have a reduced sensitivity to ABA (McCarty et al., 1991; Suzuki et al., 2003; Cao et al., 2007). Interestingly we noticed that the *vp1-1* mutant seedlings showed decreased leaf permeability compared with wild type controls (Fig. 6A). Subsequently, to investigate the possible interplay between ABA and ZmFDL1 regulated pathways, we evaluated the effect of exogenous application of the ABA hormone on the *fdl1-1* mutant. After 14 days of a treatment consisting in the application of a 1µM ABA solution (directly to the root as for the wild type; Fig. 5C) no significant reduction of cuticle dependent leaf permeability was observed, according to a chlorophyll leaching assay, in *fdl1-1* mutant seedling compared to mock-treated control (Fig. 6B).

**Figure 6.**
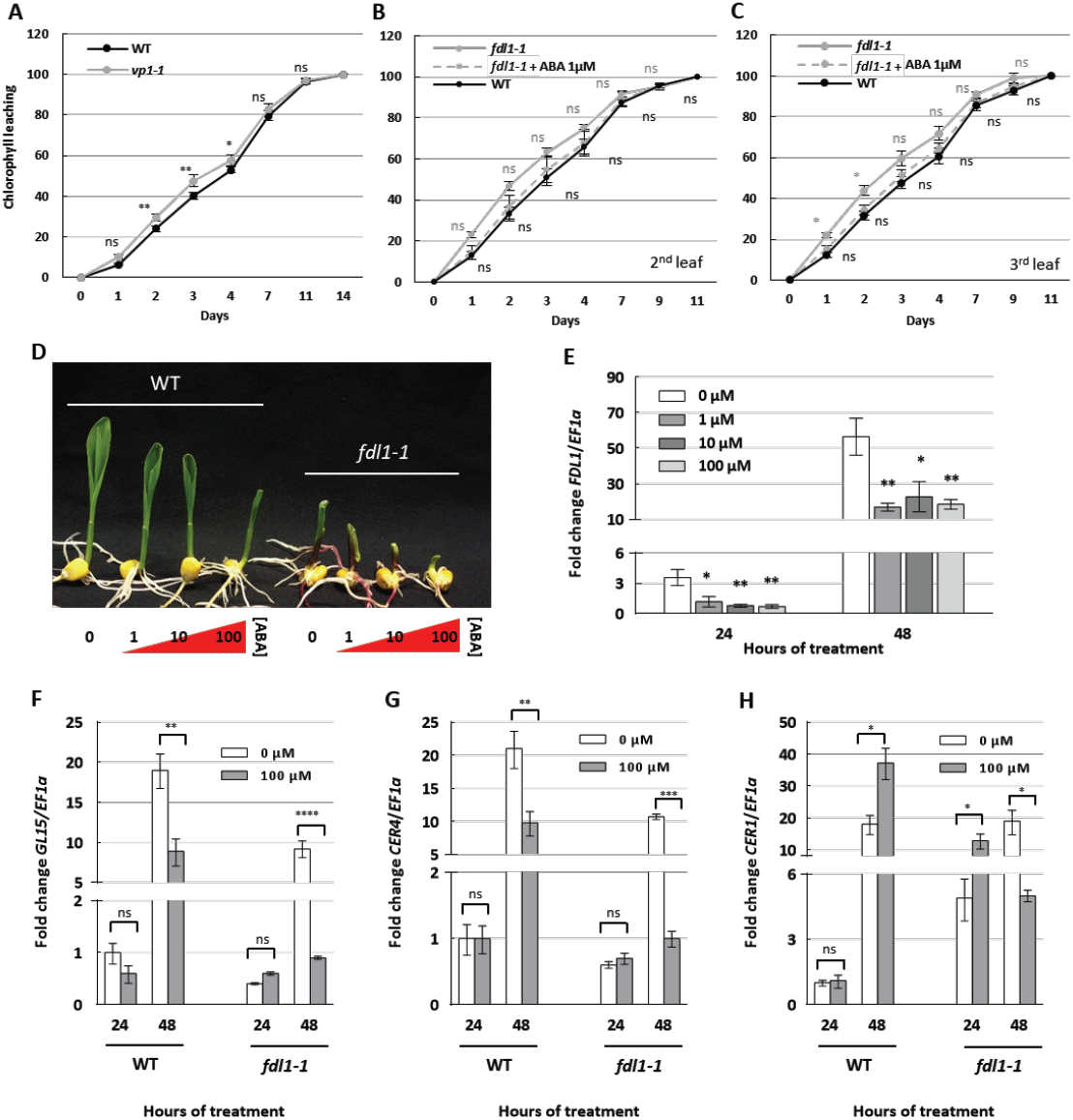
Effect of ABA treatment on leaf permeability and gene expression. The leaf permeability assessed by chlorophyll leaching assay on (A) the second fully expanded leaf of homozygous *vp1-1* and B73 wild type (WT) control plants and on the second (B) and third (C) leaves of *fdl1-1* mutant treated with control (Mock) or 1µM ABA solution and wild type (WT) plants. Values represent the mean ± SE of ten biological replicates. (D) Representative images of B73 wild type (WT) and homozygous *fdl1-1* mutant seedlings grown for 48 hours with the root apparatus into liquid solutions with ABA increasing concentration (0, 1, 10 and 100 µM). (E) Expression profile of *ZmFDL1* gene treated with control (0µM ABA) or 1, 10 and 100µM ABA solution for 24 and 48 hours. Quantitative transcript analysis of the cuticle-related (F) *ZmGL15*, (G) *ZmCER4* and (H) *ZmCER1* genes in mock (0µM ABA) or 100µM ABA treated WT and homozygous *fdl1-1* mutant seedlings for 24 and 48 hours. (D-G) Values represent the mean fold change variations ± SD of biological replicates. Comparisons were made at each time point between (A) wild type and *vp1-1* genotypes; or between (B, C) ABA and control treatments in *fdl1-1* (grey) and ABA treated *fdl1-1* mutant and wild type plants (black); or between (E-H) control and ABA treated plants. Significant differences were assessed by One-way ANOVA test or Student’s t-test (* = P < 0.05, ** = P < 0.01, *** = P < 0.001, **** = P < 0.0001, ns = not significant).

Comparisons of expression profiles in ABA and control-treated wild type plants showed a consistent down-regulation of *ZmFDL1* upon treatments with increasing concentrations of the hormone after both 24 and 48 hours of treatment (Fig. 6E). Importantly, seedlings of both *fdl1-1* and wild type control plants responded to ABA with a decrease in growth, suggesting that the perception of ABA was not altered (Fig 6D).

Expression levels of cuticle related genes were investigated in *fdl1-1* and wild type plants treated with ABA and in corresponding untreated controls. The transcript levels of both *ZmGL15* and *ZmCER4* genes were reduced upon ABA treatment in both in *fdl1-1* and wild type plants after 48 hours of ABA treatment (Fig. 6F, G). The *ZmCER1* gene (Fig. 6H) was up-regulated in *fdl1-1* after 24 hours of ABA treatment (164%) and in wild type plants after 48 hours (102%) while it was down-regulated in mutant plants after 48 hours (- 74%).

Consistent with our previous observations based on RNA-Seq data (Supplemental Table S1) and quantitative expression analysis (Fig. 3D), the *ZmGL15* and *ZmCER4* genes were down-regulated in un-treated *fdl1-1* seedlings compared to un-treated control wild type plants while *Zm*CER1 was up-regulated (Fig. 6F, G, H). The same trends were observed also upon treatment with ABA. It is remarkable that after 48 hours of ABA treatment, the reduction in expression of *ZmGL15* and *ZmCER4*, compared to their untreated controls, was more pronounced. The difference between ABA and control-treated plants was about 90% and 53% for *Zm*Gl15 (Fig. 6F), and 91% and 52% for *ZmCER4* (Fig. 6G) in *fdl1-1* mutant and wild type plants respectively. Similarly, the up-regulation of *Zm*CER1 was higher in ABA treated *fdl1-1* mutant plants (164%) than in wild type plants (102%) and was anticipated at 24 hours of treatment (Fig. 6H).

## DISCUSSION

As shown in our previous study (La Rocca et al., 2015), maize plants lacking the ZmFDL1/MYB94 transcription factor are defective in seedling development since fusions, which occur specifically in embryonic primordia and seedling organs, impair coleoptile opening as well as leaf expansion. Such fusions were attributed to the irregular deposition of intervening cuticular material between juxtaposed cell walls and to the patchy distribution of epicuticular waxes on the epidermis of the young leaves. This primary phenotypic characterization suggested that ZmFDL1 may have an important role in controlling cuticle biosynthesis in maize. This hypothesis was corroborated by the fact that AtMYB96, AtMYB94 and AtMYB30, which are its closest Arabidopsis related genes, were all shown to be involved in the regulation of cuticular wax biosynthesis (Sheperd and Wynne Griffiths, 2006; Raffaele et al., 2008; Seo et al., 2011; Lee and Suh, 2015a).

### ZmFDL1 is a key regulator of cuticle biosynthesis and deposition in maize

In the present work, a more thorough characterization, including biochemical as well as transcriptional analyses, has been undertaken to gain further insights into the role of *Zm*FDL1 in cuticle formation. The data herein obtained strongly confirm that *Zm*FDL1 is a key regulatory component of both cutin and wax biosynthesis and deposition in maize seedlings. Among cutin compounds, ω-hydroxy fatty acids and polyhydroxy-fatty acids were specifically affected in *fdl1-1* coleoptiles. In epicuticular waxes, the reduction was mainly observed in primary long chain alcohols, although a reduction of long-chain wax esters was also detected (Supplemental Fig. S1D). In agreement with the pattern observed for the morphological traits (La Rocca et al., 2015), these biochemical defects appeared transiently in germinating mutant seedlings (up to the second leaf stage), and a progressive shift to control values was observed in subsequent developmental stages for all examined compounds (Supplemental Fig. S1, S2). These data, along with the previous finding that the level of the *ZmFDL1* transcript showed a progressive decrease in the second and third leaves (La Rocca et al., 2015), provide a definitive proof that the action of *ZmFDL1* is confined to a precise developmental window delimited by the third leaf stage. This finding is in accordance with the observation that in maize, cuticle properties are different in juvenile and adult leaves. Maize juvenile leaves have a thin cuticle and are covered with epicuticular waxes crystals whereas adult leaves have a thick cuticle and an amorphous wax layer on their surfaces (Sylvester et al., 1990). Interestingly, long-chain alcohols (69%) are the main components of cuticular waxes in seedling leaves, followed by aldehydes (25%), alkanes (4%) and esters (2%) (Javelle et al., 2010). Differently, alkanes and alkyl esters are the main components in adult leaves (Bourgault et al., 2020). Cutin composition was characterized in adult leaves and shown to be mainly composed of dihydroxyhexadecanoic acid and typical members of the C18 family of cutin acids, including hydroxy and hydroxy-epoxy acids (Espelie and Kolattukudy, 1979).

The expression profile of *GLOSSY3* (*ZmGL3*), encoding another R2R3-MYB transcription factor that controls cuticle deposition, is also confined to the maize juvenile phase. Differently from *Zm*FDL1, ZmGL3, which is closely related to the Arabidopsis AtMYB60, seems specifically involved in the control of wax deposition (Liu et al., 2012). Both *ZmGL3* and *Zm*FDL1 might, in turn, interact with additional regulatory factors involved either in the maintenance of the juvenile phase or in promoting the transition from juvenile to adult phase. In this context, an interesting candidate is the *GLOSSY15* (*ZmGL15*) gene (Moose and Sisco, 1994), which encodes a transcription factor of the AP2-domain family, showing high similarity to the Arabidopsis *APETALA2* gene (*AtAP2*) (Jofuku et al., 1994) ZmGL15 is required for the maintenance of juvenile traits in the leaf epidermis and its transcriptional level is controlled by miR172 that, by down regulating *ZmGL15*, promotes the transition from juvenile to adult phase (Lauter et al., 2005).

A whole picture of the complex biosynthetic network underlying cuticle formation is proposed in figure 4 in which DEGs, as identified from our transcriptome analysis, are also reported. Lack of ZmFDL1 has an effect on the activities of genes located in different modules of the proposed pathway. With the absence of publicly available large scale data of possible *in vivo* direct targets of ZmFDL1, it is difficult to speculate whether this effect is direct or indirect. Nevertheless, we notice that lack of ZmFDL1 activity affects the expression levels of several waxes and cutin related genes, including for example transporters of cuticle components, such as the ZmGL13 wax transporter (Li et al., 2013). Interestingly, ZmGL13 was shown to be consistently down-regulation also by quantitative expression analysis (Fig. 3D).

The decrease in very long chain C32 primary alcohols (C32OH; Fig 1D) observed in mutant coleoptile waxes correlates well with the down regulation of *ZmCER4* (Fig. 3D and Supplemental Table S1), whose closest homolog is the *AtCER4* which was shown to be involved in the alcohol forming pathway (Fig. 4) (Rowland et al., 2006). Similarly, in Arabidopsis AtMYB94 activates the transcription of *AtFA3/AtCER4* and additional genes involved in wax biosynthesis (Lee and Suh, 2015b).

The *ZmWSD11* gene (Fig. 4), which corresponds to the Arabidopsis *FOLDED PETAL 1* (*AtFOP1/AtWSD11*) that encodes a bifunctional wax ester synthase/diacylglycerol acyltransferase (Takeda et al. 2013), resulted up-regulated. We also observed the up-regulation of the *ZmCER1* gene (Fig.3E), homologous to *AtCER1* that was shown to be involved in the production of very-long-chain alkanes (Bourdenx et al., 2011). However, the increased expression of *ZmCER1* in *fdl1-1* mutant did not agree with the content of these compounds observed in mutant seedlings (Supplemental Fig. S2). A general down-regulation of the genes involved in the biosynthesis of the long chain fatty acids, such as *ZmKCS17, ZmKCS22, ZmKCS24, ZmPAS2* and *ZmGL26* (Fig. 4 and Supplemental Table S1) and a strong decrease in the content of the very-long-chain aldehydes (32ALD) (Supplemental Fig. SM), precursors of the very-long-chain alkanes, could possibly explain the discrepancy between the transcript level of *ZmCER1, ZmWSD11* (Fig. 3D) and their relative products (Supplemental Fig. S2M, S1D).

The reduction of ω-hydroxy fatty acids and polyhydroxy-fatty acids (Supplemental Fig. S2I, E) was consistent with the low expression of genes involved in the biosynthesis of cutin precursors and cutin monomers, such as the *ZmONI3, ZmHTH1, ZmCUS2* and *ZmAAE16, ZmCER8, ZmLACS2* and *ZmLACS4* (Fig. 4 and Supplemental Table S1).

Interestingly, in rice, OsONI3, showing high level of similarity with an Arabidopsis HOTHEAD (HTH) protein (Akiba et al., 2014), was suggested to encode an omega-hydroxy dehydrogenase catalysing the biosynthesis of the long chain α, ω-dicarboxylic fatty acids (Yephremov et al., 1999). Interestingly, the seedling morphology of the *OsONI3* mutant highly resembles that of *fdl1-1*, showing regions of fusion involving embryo leaf primordia as well as shoot leaves. This may suggest that the proper synthesis of cutin related compounds is required to prevent post-germinative adhesion among organs.

The results of a recent genome wide analysis, based on DAP-Seq (Liu et al., 2020), identified 6 of our selected genes as possible direct targets of ZmFDL1 (p-value hypergeometric 1.6E-07). These direct targets are Zm00001d004125 (*ZmACC2*); Zm00001d044579 (*ZmKCS22*/*ZmCER60*); Zm00001d039053 (ZmKCS15); Zm00001d053127 (*ZmLACS2*); Zm00001d017251 (Zm*CER1*) and Zm00001d043853 (Zm*WSD11*). This indicates the robustness of our approach.

Furthermore, the strong enrichment of down-regulated DEGs in the KEGG “zma04712: Circadian rhythm” pathway (Supplemental Table S2) could reflect the conserved function between ZmFDL1 and its Arabidopsis homolog AtMYB96, which was shown to be connected with the clock, to shape the circadian gating of abscisic acid (ABA) responses (Lee et al., 2016).

### *ZmFDL1* and associated cuticle-related genes mediate leaf permeability under drought and ABA treatments

We have shown in the present study that ZmFDL1, besides promoting organ separation, is required in maize seedling leaves to control cuticle-dependent leaf permeability, to reduce the trans-cuticular water loss and eventually to preserve the leaf water status (Fig. 2). Several studies have shown that cuticle formation responds to environmental conditions, suggesting that the cuticle retains an active role in the plant response to environmental stress conditions, among them water scarcity (Xue et al., 2017). Our study provides support to these findings and further indicates that in maize seedlings this response is mediated by ZmFDL1. It also provides evidence that additional regulators such as ZmGL15, as well the ABA hormone signalling pathway, are involved. Cuticle-dependent leaf permeability, as measured through chlorophyll leaching analysis on B73 maize seedlings, decreased under limited water supply as well as through the exogenous administration of the ABA hormone (Fig. 5A, B). Moreover, the application of a drought stress treatment (LW) to developing seedlings caused a change in the expression profile of *ZmFDL1* and associated genes (Fig. 5, 7, and Supplemental Fig. S4).

**Figure 7.**
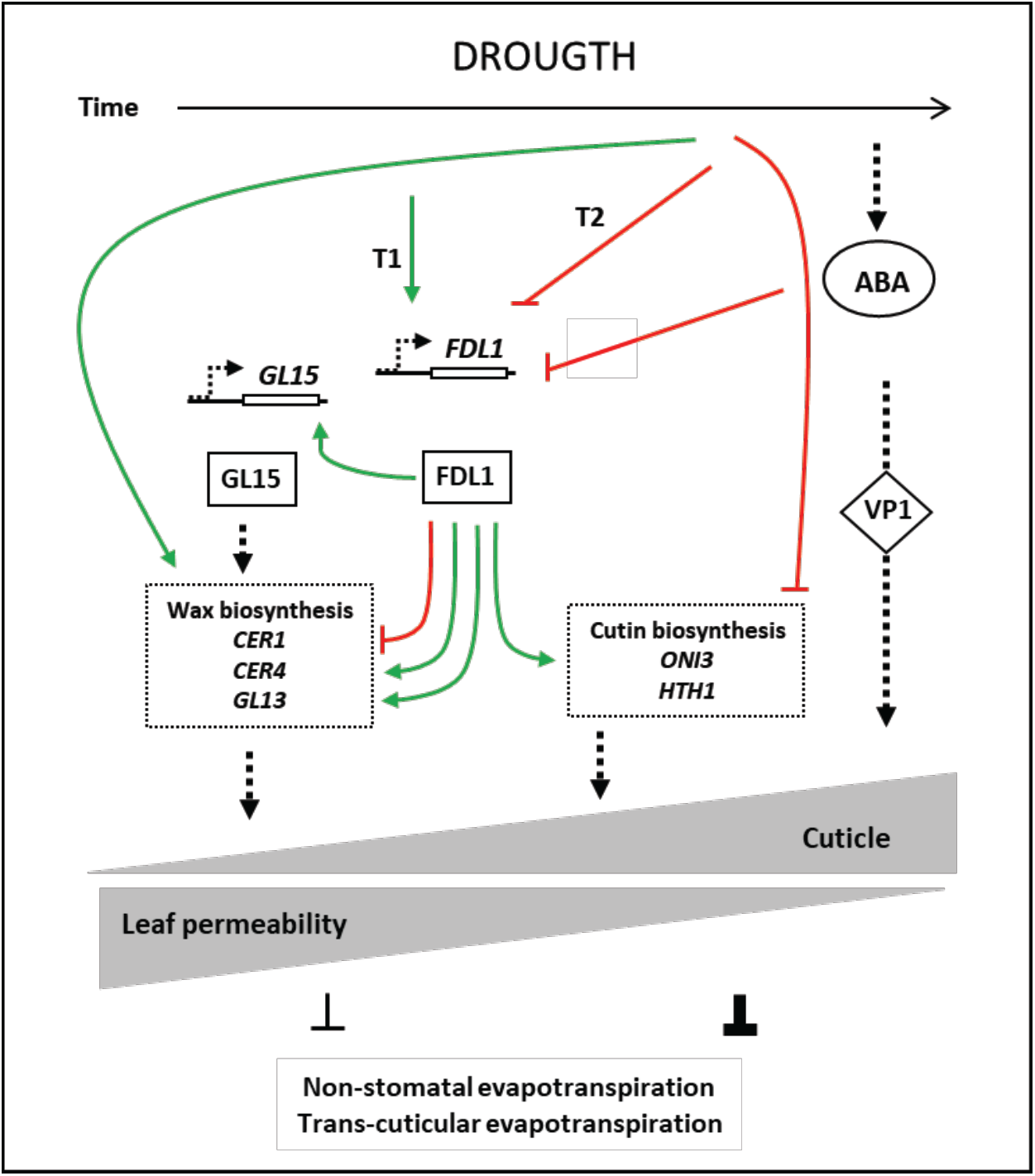
The proposed roles of ZmFDL1 and ABA in the regulation of cuticle-dependent leaf permeability under drought. The green arrows indicate transcriptional activation and the T red lines indicate transcriptional repression. The black dotted arrows indicate activation but the mode of interaction is not known.

Under limited watering conditions (LW), both *ZmFDL1* and *ZmGL15* showed an early peak of expression with respect to normally watered plants (NW). This observation applies also to *ZmCER4*, while the expression profiles of both *ZmCER1, ZmWSD11* and *ZmGL13* showed a neat peak that was not observed in NW condition. We may speculate that the effect of drought on the expression profile of *ZmCER4* is modulated to a greater extent by the positive action exerted by ZmFDL1, which might also mediate the positive effect exerted by drought on the *ZmGL13* transcript level. Differently, if we assume from our previous data that ZmFDL1 is a repressor of *ZmCER1* and *ZmWSD11*, their up-regulation observed under drought might be independent from the action of this MYB factor. Overall our data indicate that genes involved in waxes biosynthesis (*ZmCER1, Zm*CER4 and *Zm*WSD11), and eventually the total amount of cuticular waxes, are increased in response to drought stress conditions, thus improving plant protection from water loss. This finding is in agreement with a number of studies conducted for wax-related *Arabidopsis* genes. One example is constituted by *AtCER1*, that controls the biosynthesis of very-long-chain-alkane (Bernard et al., 2012) and was correlated with drought stress responses (Bourdenx et al., 2011). Its overexpression in plants significantly increases the production of alkanes and leads to increased drought tolerance (Bourdenx et al., 2011). Moreover, the expression of *AtWAX2*, which is involved in both cutin and wax biosynthesis, was shown to be induced by drought, ABA, low temperature and salinity in cucumber (Chen et al., 2003).

Our analysis suggests that *ZmFDL1* and waxes-related genes, through increasing their transcriptional activity, provide a rapid response to the drought stimulus. This response seems transient, since at later stages, when the stress condition is more severe, as indicated by the high expression level of the *ZmRAB18*, their transcript levels are diminished. A negative feedback might be produced, which causes the repression of wax biosynthesis. The observed gene pattern might also reflect the pattern of leaf growth meaning that in leaves approaching an advanced developmental stage, cuticle biosynthesis is diminished.

The role of ABA, in mediating the response to water deficit has been known since many years (Bartels and Sunkar, 2005; Shinozaki and Yamaguchi-Shinozaki, 2007) and the effect of ABA treatment on cuticle composition and related gene expression has been reported in Arabidopsis (Kosma et al., 2009) and tomato (Martin et al., 2017). We observed in the present study that absence of ABA perception in *vp1* mutant seedlings resulted in increased cuticle-dependent leaf permeability (Fig. 7), while a slight decrease in leaf permeability was obtained through the application of exogenous ABA treatment (Fig. 7), thus showing, for the first time in maize, the involvement of ABA in this process.

We also observed that ABA exogenous application to B73 wild type seedlings caused severe repression of *ZmFDL1*, after both 24 and 48 hours of treatment. As expected, the repression, at the transcriptional level, was also observed for both *ZmGL15* and *ZmCER4*, although only after 48 hours of ABA treatment (Fig 6F, G). A different pattern was observed for *ZmCER1* that was up-regulated in the ABA treated plants (Fig. 6E). The early response observed in the *fdl1-1* mutant plants might be due to the absence of the repressive role exerted by *Zm*FDL1 on the transcription of this gene. Accordingly, in Arabidopsis, the level of waxes and in particular of alkanes, were shown to increase following ABA application (Kosma et al., 2009).

The unexpected reduction of the expression of cuticle related genes observed in this experiment, which resembles what observed at later stages in seedlings grown under limited watering (Fig.5E-J, Days 5,7) might be due the high concentrations of the exogenous ABA applied, which are considerably higher than physiological levels. These experimental conditions, with increasing ABA concentration are plausibly mimicking severe drought stress (Dallmier and Stewart, 1992). It is remarkable that the change in gene expression of *ZmGL15* and *ZmCER4* caused by ABA treatment, although qualitatively similar in both wild type and *fdl1-1* mutant plants, was considerably more pronounced in the latter. Taken together the data from the expression analysis strongly suggest the existence of two signalling pathways promoted by the ABA hormone for the regulation of cuticular wax biosynthesis, of which one is independent and a second requires the action of ZmFDL1. The two pathways may act additively in controlling the transcription of biosynthetic genes.

In summary (Fig. 7), we showed in this study that the *Zm*FDL1 transcription factor is a key regulator of cuticle deposition and mediates the active response of cuticle to water scarcity condition. *Zm*FDL1 sensitivity to drought stress, consisting in the up-regulation of its transcript, occurred very early during seedling leaf development, as observed in the second leaf not yet emerged (Fig. 5G). This early *Zm*FDL1 response, accompanied by the activation of the regulatory gene *ZmGL15*, may allow plants to rapidly cope with water scarcity conditions in young tissues, by stimulating the activity of genes involved in cuticle biosynthesis, and more specifically in the wax deposition. A reduction in non-stomatal evapotranspiration is thus achieved that would prevent water deficit. It appeared, however, that the FDL1 mediated response is less effective at later developmental stages as well as when the severity of the imposed stress is high. We also showed that in maize seedlings ABA perception (signalling) influences cuticle dependent leaf-permeability and ABA has a negative effect on *Zm*FDL1 transcription. Although mechanisms of interaction between the hormone signalling pathway and the *Zm*FDL1 regulatory pathway remains to be elucidated, our model suggests that the two pathways interact to modulate the expression of cuticle related genes.

## MATERIALS AND METHODS

### Plant materials and growth conditions

The maize (*Zea mays*) *fdl1-1* mutant, first described by La Rocca *et al* (2015)., was introgressed into B73, H99 and Mo17 inbred lines. In all the experiments performed, homozygous mutants and their wild type control plants were from the same F_2_ segregating progeny. Plants were grown under long day photoperiod (16 h of light/8 h of dark) in a growth chamber with controlled temperature (25 °C night, 30°C day) and with photon fluence of 70 µmol m^-2^ s^-1^ in pots containing S.Q.10 substrate (peat, sand, compost) (Vigorplant).

For drought treatment, maize seedlings were germinated and grown in soil under normal-watered (NW) condition (RSWC=80%) until 5 days after germination (DAS) and the next day (6 DAS, day 0) the drought stress was imposed by withholding irrigation. To monitor the drought stress level, the relative soil water contents (RSWC) were measured (Janeczko *et at.* 2016), as previously described (Castorina et al., 2018). To keep the plants under NW or low-watered (LW) conditions the RSWC was measured every day and the pots were watered as necessary until the end of the experiments. For the cuticle-dependent epidermal permeability assay the drought (RSWC=40%) was imposed up to 18 DAS (Fig. 2D) and for each treatment 7 pots were used. Whereas, for the expression analysis of cuticle related genes the drought (RSWC=30%) was imposed up to 13 DAS (Fig. 6 and Supplementary Figure S4) and for each time point the second leaf from four/five independent plants per treatment were sampled.

To mimic drought stress, plants were treated with exogenous applications of the abscisic acid (ABA) hormone (Duchefa) as previously described by (Riboni et al., 2016). The *fdl1-1* and B73 wild type seedlings at the coleoptile developmental stage has been treated with 1ml of ABA (1 µM) or mock control (0.001% v/v ethanol) solution. Starting from 4 DAS, the treatments have been dispensed, every other day, watering directly the roots up to 14 DAS. For each treatment (Mock and ABA) the second leaf, from a minimum of 7 independent plants per genotype, was taken (Fig. 2E, Supplementary Figure S3).

To analyse the transcript regulation effect of the ABA on *ZmFDL1* and other cuticle-related genes, B73 wild type and *fdl1-1* seedlings have been treated dipping only the root apparatus into an ABA (1, 10 and 100 µM) or mock control solution. After 24 and 48 hours of treatment, only the green tissues (coleoptile and leaves) from a minimum of six independent plants per genotype have been sampled for subsequent total RNA extraction.

### Cuticular Cutin and Wax Analysis

Cuticle composition was analysed in *fdl1-1* and wild type seedlings at succeeding developmental stages: coleoptile stages, first emerging leaf, second emerging leaf and third emerging leaf at 2, 6, 10 and 14 days after sowing, respectively. Cutin and epicuticular waxes were extracted and identified by the combination of gas chromatography on column injection (GC-OCI) and gas chromatography-mass spectrometry (GC-MS) performed according to previously described methods (Domergue et al., 2010; Bourdenx et al., 2011).

### Chlorophyll leaching assay, leaf temperature and water loss

For the chlorophyll leaching analysis, second, third and fourth fully expanded leaves were taken from a minimum of five independent plants per genotype and dissected into pieces of 8 cm in length as measured from the apex. Leaf sectors were weighed, immersed in ethanol 80% and incubated in the dark at room temperature. Absorbance was measured every day at 647 and 664 nm with a spectrophotometer (Agilent Technologies Cary 60 UV-Vis) to quantify the chlorophyll released in the solution and measurements were performed until complete chlorophyll extraction. The micromolar concentration of chlorophyll was calculated using the equation: total micromoles chlorophyll = 7.93 x A664 + 19.53 x A647 (Lolle et al., 1997). The data obtained were normalized per gram of fresh weight and area of the 8 cm long leaf pieces to be expressed as a percentage of the total chlorophyll.

Thermal images of second fully expanded leaves were taken from 14 old-days *fdl1-1* and wild type seedlings with a semi-automated long-wave infrared cameras system (FLIR T650sc). The temperature of the leaves was measured using the FLIR ResearchIR Max software.

To determine the time course of the seedling water loss, 10-old-days seedlings were detached and weighed immediately. Seedling weight was then estimated at designated time intervals and water loss was calculated as the percentage of fresh weight (FW) based on the initial weight. Several biological replicates were measured for each genotype.

### Cell density and stomatal index analysis

To measure stomatal density and stomatal index, a leaf surface imprint method was used as previously described (Castorina et al., 2018). We analysed second, third and fourth fully expanded leaves of *fdl1-1* mutant and wild type plants. Stomatal index (SI) was determined as [number of stomata / (number of epidermal cells + number of stomata)] × 100. One-way ANOVA test was performed with the statistical package SPSS 21.0.

### RNA, cDNA Preparation and quantitative gene expression analysis

Total RNA was extracted from maize tissues using the TRIzol Reagent (Life Technologies) and treated with RQ1 RNase-Free DNase (Promega) according to the manufacturer’s instructions. First-strand complementary DNA (cDNA) was synthetized with the SuperScript® III First-Strand Synthesis System (Invitrogen) from 1000 ng of total RNA, according to the manufacturer’s instructions. Real-Time PCR was performed with the 7300 Real-Time PCR System (Applied Biosystems), using GoTaq qPCR Master Mix (Promega), in a final volume of 10 μl. The following cycle was used: 10 min pre-incubation at 95°C, followed by 40 cycles of 15 s at 95°C and 1 min at 60°C. The relative transcript level of each gene was calculated by the 2^-ΔΔCt^ method (Livak and Schmittgen, 2001) using the expression of the *ZmEF1a* gene as a reference. The gene-specific primers are listed in Supplementary Table S3.

The RNA extractions and purifications for RNA sequencing were performed on seedlings at coleoptile stage of *fdl1-1* and wild type plants 2 days after germination collecting 3 plants/genotypes (n=3) and homogenized in liquid nitrogen. The total RNA extraction was performed with the Pure Link RNA Mini Kit (Invitrogen). 4 μg of total RNA with an RNA Integrity Number (RIN) ≥ 8 (Bioanalyzer 2100, Agilent Technologies, Santa Clara, CA, United States) were sent to the IGA Technology Services4 (Udine, Italy) for library preparation (TruSeq Stranded mRNA, Illumina) and sequencing on HiSeq 2000 platform (single-read 50 bp, 6-plex, about 20 million reads/sample).

### Differential expression analysis and functional enrichment analyses

Reads were mapped on the Zm00001d.2 gene models annotation of the B73 reference assembly (ZmB73_RefGen_v4) of the maize genome, as obtained from http://plants.ensembl.org/info/website/ftp/index.html, using the bowtie2 program (Langmead and Salzberg, 2012).

Estimation of gene expression levels was performed using RSEM (Li and Dewey, 2011). Differential expression analysis was performed applying the latest versions of DESeq2 (Love et al., 2014) and Limma (Ritchie et al., 2015) to RSEM estimated reads counts. Only genes with a median read count of 10 or more were considered (10 reads in at least 3 samples). Genes showing a false discovery rate lower than 0.05 according to both tools were considered differentially expressed. Gene ontology annotation of maize genes according to (Wimalanathan et al., 2018) was obtained from http://datacommons.cyverse.org/browse/iplant/home/shared/commons_repo/curated/Carolyn_Lawrence_Dill_maize-GAMER_July_2017_V.1. Functional enrichment analyses were performed by means of a custom script implementing a Fisher exact test and the Benjamini Hochberg procedure for the correction of multiple testing.

Graphical representation of the results of Gene Ontology functional enrichment was performed by means of the REViGO tool (Supek et al., 2011), as available from http://revigo.irb.hr/, using default parameters. Gene ontology terms of interest were highlighted manually.

Annotation of maize metabolic pathways according to the Mercator MapMan ontology (Schwacke et al., 2019) was obtained from https://mapman.gabipd.org/mapmanstore. Functional enrichment of metabolic pathways was performed by using MapMan version 3.5.1.

Publicly available KEGG pathways for Z. mays were obtained directly from the KEGG PATHWAY repository, as listed from <https://www.genome.jp/dbget-bin/get_linkdb?-t+pathway+gn:T01088.> The database was accessed on March 10th 2020.

## ACKNOWLEDGEMENTS

We acknowledge the Maize Genetics Cooperation Stock Center, Urbana, IL, USA for providing the *vp1* mutant.

## LITERATURE CITED

Aharoni A, Dixit S, Jetter R, Thoenes E, van Arkel G, Pereira A (2004) The SHINE clade of AP2 domain transcription factors activates wax biosynthesis, alters cuticle properties, and confers drought tolerance when overexpressed in Arabidopsis. Plant Cell 16: 2463–2480

Akiba T, Hibara KI, Kimura F, Tsuda K, Shibata K, Ishibashi M, Moriya C, Nakagawa K, Kurata N, Itoh JI, et al (2014) Organ fusion and defective shoot development in *oni3* mutants of rice. Plant Cell Physiol 55: 42–51

Bartels D, Sunkar R (2005) Drought and Salt Tolerance in Plants. CRC Crit Rev Plant Sci 24: 23–58

Bernard A, Domergue F, Pascal S, Jetter R, Renne C, Faure JD, Haslam RP, Napier JA, Lessire R, Joubès J (2012) Reconstitution of plant alkane biosynthesis in yeast demonstrates that Arabidopsis ECERIFERUM1 and ECERIFERUM3 are core components of a very-long-chain alkane synthesis complex. Plant Cell 24: 3106–3118

Bernard A, Joubès J (2013) Arabidopsis cuticular waxes: Advances in synthesis, export and regulation. Prog Lipid Res 52: 110–129

Bi H, Luang S, Li Y, Bazanova N, Morran S, Song Z, Perera MA, Hrmova M, Borisjuk N, Lopato S (2016) Identification and characterization of wheat drought-responsive MYB transcription factors involved in the regulation of cuticle biosynthesis. J Exp Bot 67: 5363–5380

Bourdenx B, Bernard A, Domergue F, Pascal S, Léger A, Roby D, Pervent M, Vile D, Haslam RP, Napier JA, et al (2011) Overexpression of Arabidopsis *ECERIFERUM1* promotes wax very-long-chain alkane biosynthesis and influences plant response to biotic and abiotic stresses. Plant Physiol 156: 29–45

Bourgault R, Matschi S, Vasquez M, Qiao P, Sonntag A, Charlebois C, Mohammadi M, Scanlon MJ, Smith LG, Molina I (2020) Constructing functional cuticles: analysis of relationships between cuticle lipid composition, ultrastructure and water barrier function in developing adult maize leaves. Ann Bot 125: 79–91

Burow GB, Franks CD, Xin Z (2008) Genetic and physiological analysis of an irradiated bloomless mutant (epicuticular wax mutant) of sorghum. Crop Sci 48: 41–48

Cao X, Costa LM, Biderre-Petit C, Kbhaya B, Dey N, Perez P, McCarty DR, Gutierrez-Marcos JF, Becraft PW (2007) Abscisic acid and stress signals induce *Viviparous1* expression in seed and vegetative tissues of maize. Plant Physiol 143: 720–731

Castorina G, Persico M, Zilio M, Sangiorgio S, Carabelli L, Consonni G (2018) The maize *lilliputian1 (lil1*) gene, encoding a brassinosteroid cytochrome P450 C-6 oxidase, is involved in plant growth and drought response. Ann Bot 122: 227–238

Chen X, Goodwin SM, Boroff VL, Liu X, Jenks MA (2003) Cloning and characterization of the *WAX2* gene of Arabidopsis involved in cuticle membrane and wax production. Plant Cell 15: 1170–1185

Cui F, Brosché M, Lehtonen MTT, Amiryousefi A, Xu E, Punkkinen M, Valkonen JPPT, Fujii H, Overmyer K (2016) Dissecting Abscisic Acid Signaling Pathways Involved in Cuticle Formation. Mol Plant 9: 926–938

Dallmier KA, Stewart CR (1992) Effect of exogenous abscisic acid on proline dehydrogenase activity in maize (*Zea mays L.*). Plant Physiol 99: 762–764

Domergue F, Vishwanath SJ, Joubès J, Ono J, Lee JA, Bourdon M, Alhattab R, Lowe C, Pascal S, Lessire R, et al (2010) Three Arabidopsis fatty Acyl-coenzyme a reductases, FAR1, FAR4, and FAR5, generate primary fatty alcohols associated with suberin deposition. Plant Physiol 153: 1539–1554

Espelie KE, Kolattukudy PE (1979) Composition of the aliphatic component of ‘suberin’ from the bundle sheaths of *Zea mays* leaves. Plant Sci Lett 15: 225–230

Fich EA, Segerson NA, Rose JKC (2016) The Plant Polyester Cutin: Biosynthesis, Structure, and Biological Roles. Annu Rev Plant Biol 67: 207–233

Giraudat J, Hauge BM, Valon C, Smalle J, Parcy F, Goodman HM (1992) Isolation of the Arabidopsis *ABI3* gene by positional cloning. Plant Cell 4: 1251–1261

Haslam TM, Kunst L (2013) Extending The Story Of Very-Long-Chain Fatty Acid Elongation. Plant Sci 210: 93–107

Ingram G, Nawrath C (2017) The roles of the cuticle in plant development: organ adhesions and beyond. J Exp Bot 68: 5307–5321

Javelle M, Vernoud V, Depege-Fargeix N, Arnould C, Oursel D, Domergue F, Sarda X, Rogowsky PM (2010) Overexpression of the Epidermis-Specific Homeodomain-Leucine Zipper IV Transcription Factor OUTER CELL LAYER1 in Maize Identifies Target Genes Involved in Lipid Metabolism and Cuticle Biosynthesis. Plant Physiol 154: 273–286

Jiao Y, Peluso P, Shi J, Liang T, Stitzer MC, Wang B, Campbell MS, Stein JC, Wei X, Chin CS, et al (2017) Improved maize reference genome with single-molecule technologies. Nature 546: 524–527

Jofuku KD, den Boer BGW, Van Montagu M, Okamuro JK (1994) Control of Arabidopsis flower and seed development by the homeotic gene *APETALA2*. Plant Cell 6: 1211–1225

Kosma DK, Bourdenx B, Bernard A, Parsons EP, Lü S, Joubès J, Jenks MA (2009) The impact of water deficiency on leaf cuticle lipids of Arabidopsis. Plant Physiol 151: 1918–1929

Langmead B, Salzberg SL (2012) Fast gapped-read alignment with Bowtie 2. Nat Methods 9: 357–359

Lauter N, Kampani A, Carlson S, Goebel M, Moose SP (2005) microRNA172 down-regulates *glossy15* to promote vegetative phase change in maize. Proc Natl Acad Sci U S A 102: 9412–7

Lee HG, Mas P, Seo PJ (2016) MYB96 shapes the circadian gating of ABA signaling in Arabidopsis. Sci Rep 6: 1–11

Lee SB, Suh MC (2015a) Cuticular wax biosynthesis is up-regulated by the MYB94 transcription factor in arabidopsis. Plant Cell Physiol 56: 48–60

Lee SB, Suh MC (2015b) Advances in the understanding of cuticular waxes in Arabidopsis thaliana and crop species. Plant Cell Rep 34: 557–572

Li-Beisson Y, Shorrosh B, Beisson F, Andersson MX, Arondel V, Bates PD, Baud S, Bird D, DeBono A, Durrett TP, et al (2013) Acyl-Lipid Metabolism. Arab B 11: e0161

Li B, Dewey CN (2011) RSEM: accurate transcript quantification from RNA-Seq data with or without a reference genome. BMC Bioinformatics 12: 323

Li F, Wu X, Lam P, Bird D, Zheng H, Samuels L, Jetter R, Kunst L (2008) Identification of the wax ester synthase/acyl-coenzyme a:diacylglycerol acyltransferase WSD1 required for stem wax ester biosynthesis in Arabidopsis. Plant Physiol 148: 97–107

Li L, Du Y, He C, Dietrich CR, Li J, Ma X, Wang R, Liu Q, Liu S, Wang G, et al (2019) Maize *glossy6* is involved in cuticular wax deposition and drought tolerance. J Exp Bot 70: 3089–3099

Li L, Li D, Liu S, Ma X, Dietrich CR, Hu HC, Zhang G, Liu Z, Zheng J, Wang G, et al (2013) The Maize *glossy13* gene, cloned via BSR-Seq and Seq-Walking encodes a putative ABC transporter required for the normal accumulation of epicuticular waxes. PLoS One 8: 1–13

Liu S, Yeh CT, Tang HM, Nettleton D, Schnable PS (2012) Gene mapping via bulked segregant RNA-Seq (BSR-Seq). PLoS One 7: 1–8

Liu X, Bourgault R, Strable J, Galli M, Chen Z, Dong J, Molina I, Gallavotti A (2020) the Fused Leaves1/Adherent1 Regulatory Module Is Required for Maize Cuticle Development and Organ Separation. 1–46

Livak KJ, Schmittgen TD (2001) Analysis of relative gene expression data using real-time quantitative PCR and the 2-ΔΔCT method. Methods 25: 402–408

Lolle SJ, Berlyn GP, Engstrom EM, Krolikowski KA, Reiter W, Pruitt RE (1997) Developmental Regulation of Cell Interactions in the Arabidopsis fiddlehead-1 Mutant : A Role for the Epidermal Cell Wall and Cuticle. 321: 311–321

Love MI, Huber W, Anders S (2014) Moderated estimation of fold change and dispersion for RNA-seq data with DESeq2. Genome Biol 15: 550

Mao H, Wang H, Liu S, Li Z, Yang X, Yan J, Li J, Tran LSP, Qin F (2015) A transposable element in a NAC gene is associated with drought tolerance in maize seedlings. Nat Commun 6: 1–7

Martin LBB, Romero P, Fich EA, Domozych DS, Rose JKC (2017) Cuticle biosynthesis in tomato leaves is developmentally regulated by abscisic acid. Plant Physiol 174: 1384–1398

McCarty DR, Hattori T, Carson CB, Vasil V, Lazar M, Vasil IK (1991) The *Viviparous-1* developmental gene of maize encodes a novel transcriptional activator. Cell 66: 895–905

Moose SP, Sisco PH (1994) Glossy15 Controls the Epidermal Juvenile-to-Adult Phase Transition in Maize. Plant Cell 6: 1343–1355

Moriya Y, Itoh M, Okuda S, Yoshizawa AC, Kanehisa M (2007) KAAS: An automatic genome annotation and pathway reconstruction server. Nucleic Acids Res 35: 182–185

Okuda S, Yamada T, Hamajima M, Itoh M, Katayama T, Bork P, Goto S, Kanehisa M (2008) KEGG Atlas mapping for global analysis of metabolic pathways. Nucleic Acids Res 36: 423–426

Park JJ, Jin P, Yoon J, Yang J Il, Jeong HJ, Ranathunge K, Schreiber L, Franke R, Lee IJ, An G (2010) Mutation in *Wilted Dwarf and Lethal 1 (WDL1*) causes abnormal cuticle formation and rapid water loss in rice. Plant Mol Biol 74: 91–103

Raffaele S, Vailleau F, Leger A, Joubes J, Miersch O, Huard C, Blee E, Mongrand S, Domergue F, Roby D (2008) A MYB transcription factor regulates very-long-chain fatty acid biosynthesis for activation of the hypersensitive cell death response in Arabidopsis. Plant Cell 20: 752–767

Riboni M, Test AR, Galbiati M, Tonelli C, Conti L (2016) ABA-dependent control of GIGANTEA signalling enables drought escape via up-regulation of FLOWERING LOCUS T in *Arabidopsis thaliana*. J Exp Bot 67: 6309–6322

Ritchie ME, Phipson B, Wu D, Hu Y, Law CW, Shi W, Smyth GK (2015) Limma powers differential expression analyses for RNA-sequencing and microarray studies. Nucleic Acids Res 43: e47

La Rocca N, Manzotti PS, Cavaiuolo M, Barbante A, Vecchia FD, Gabotti D, Gendrot G, Horner DS, Krstajic J, Persico M, et al (2015) The maize *fused leaves1 (fdl1*) gene controls organ separation in the embryo and seedling shoot and promotes coleoptile opening. J Exp Bot 66: 5753–5767

Rowland O, Zheng H, Hepworth SR, Lam P, Jetter R, Kunst L (2006) *CER4* encodes an alcohol-forming fatty acyl-coenzyme A reductase involved in cuticular wax production in Arabidopsis. Plant Physiol 142: 866–877

Schwacke R, Ponce-Soto GY, Krause K, Bolger AM, Arsova B, Hallab A, Gruden K, Stitt M, Bolger ME, Usadel B (2019) MapMan4: A Refined Protein Classification and Annotation Framework Applicable to Multi-Omics Data Analysis. Mol Plant 12: 879–892

Seo PJ, Lee SB, Suh MC, Park M-J, Go YS, Park C-M (2011) The MYB96 transcription factor regulates cuticular wax biosynthesis under drought conditions in Arabidopsis. Plant Cell 23: 1138–1152

Seo PJ, Xiang F, Qiao M, Park JY, Lee YN, Kim SG, Lee YH, Park WJ, Park CM (2009) The MYB96 transcription factor mediates abscisic acid signaling during drought stress response in Arabidopsis. Plant Physiol 151: 275–289

Sheperd T, Wynne Griffiths D G (2006) The effects of stress on plant cuticular\rwaxes. New Phytol 171: 469–499

Shinozaki K, Yamaguchi-Shinozaki K (2007) Gene networks involved in drought stress response and tolerance. J Exp Bot 58: 221–227

Supek F, Bošnjak M, Škunca N, Šmuc T (2011) REVIGO Summarizes and Visualizes Long Lists of Gene Ontology Terms. PLoS One 6: e21800

Suzuki M, Ketterling MG, Li QB, McCarty DR (2003) Viviparous1 alters global gene expression patterns through regulation of abscisic acid signaling. Plant Physiol 132: 1664–1677

Sylvester AW, Cande WZ, Freeling M (1990) Division and differentiation during normal and liguleless-1 maize leaf development. Development 110: 985–1000

Vogg G, Fischer S, Leide J, Emmanuel E, Jetter R, Levy AA, Riederer M (2004) Tomato fruit cuticular waxes and their effects on transpiration barrier properties: Functional characterization of a mutant deficient in a very-long-chain fatty acid β-ketoacyl-CoA synthase. J Exp Bot 55: 1401–1410

Wimalanathan K, Friedberg I, Andorf CM, Lawrence-Dill CJ (2018) Maize G. Annotation—Methods, Evaluation, and Review (maize-GAMER). Plant Direct 2: 1–15

Xue D, Zhang X, Lu X, Chen G, Chen ZH (2017) Molecular and evolutionary mechanisms of cuticular wax for plant drought tolerance. Front Plant Sci 8: 1–12

Yeats TH, Rose JKC (2013) The formation and function of plant cuticles. Plant Physiol 163: 5–20

Yephremov A, Wisman E, Huijser P, Huijser C, Wellesen K, Saedler H (1999) Characterization of the *FIDDLEHEAD* gene of Arabidopsis reveals a link between adhesion response and cell differentiation in the epidermis. Plant Cell 11: 2187–2201

Zhou X, Li L, Xiang J, Gao G, Xu F, Liu A, Zhang X, Peng Y, Chen X, Wan X (2015) *OsGL1-3* is involved in cuticular wax biosynthesis and tolerance to water deficit in rice. PLoS One 10: 1–18

