## Supplemental Figures and Tables for "The drought-responsive *ZmFDL1* gene regulates cuticle biosynthesis and cuticle-dependent leaf permeability"

**Short Title**
ZmFDL1 regulates cuticle-dependent leaf permeability.

**One-Sentence Summary**
Cuticle biosynthesis and cuticle-mediated drought-response during the juvenile phase of maize plant growth, are regulated by the MYB transcription factor fused leaves1 (ZmFDL1) and influenced by ABA.

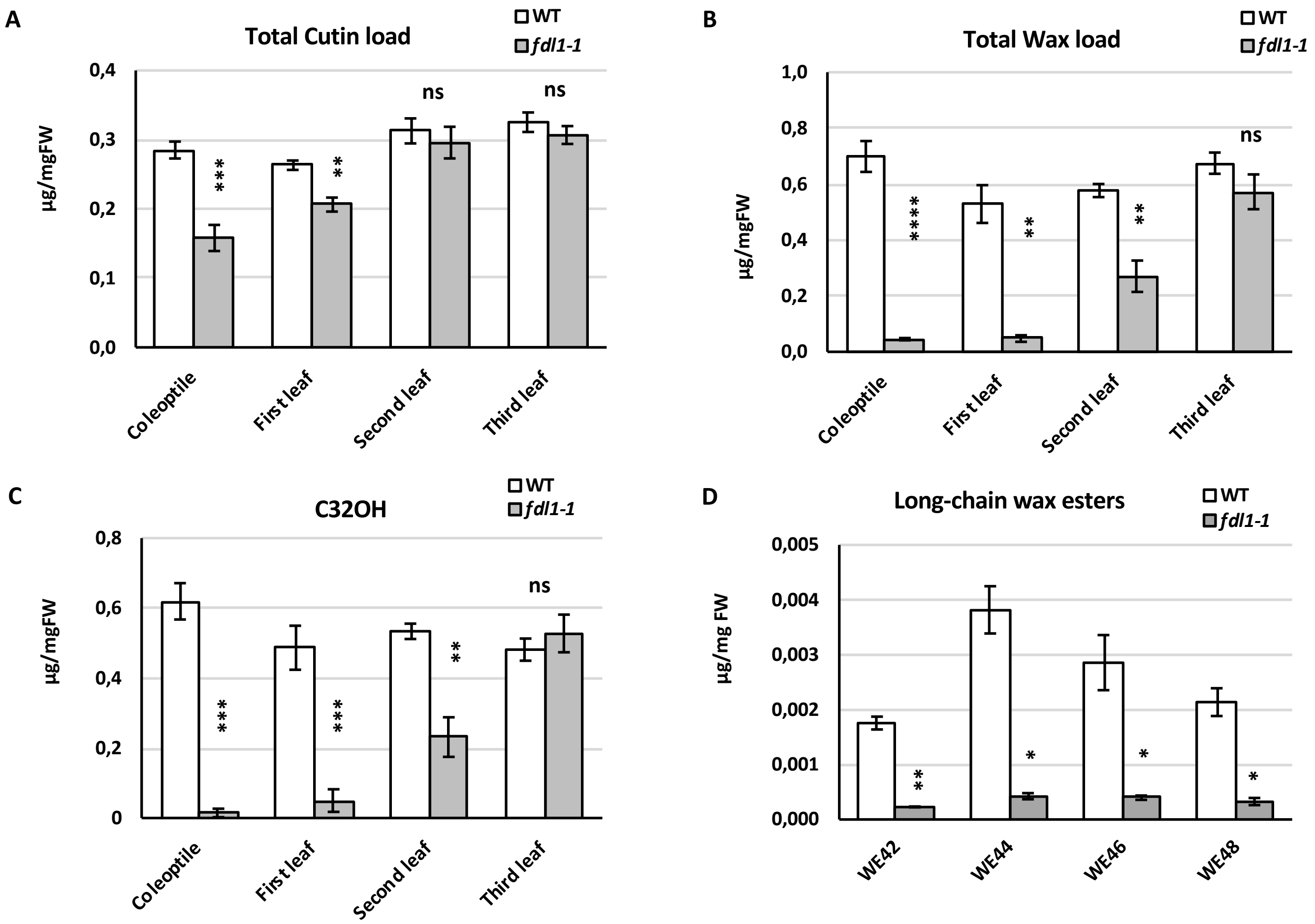


**Supplemental Figure S1. Developmental phase-dependent cuticle composition.** Total cutin (A) and wax (B) loads. (C) Relative amounts of 32C long chain primary alcohol (C32OH) at coleoptile, first leaf, second leaf and third leaf developmental stages in wild type (WT) and homozygous mutant plants (*fdl1-1*). (D) Relative amounts of the wax esters (WE) in *fdl1-1* mutant and wild type (WT) seedlings at coleoptile developmental stage. Values represent the mean ± SE of five biological replicates. Significant differences were assessed by Student's t-test (* = P < 0.05, ** = P < 0.01, *** = P < 0.001, **** = P < 0.0001, ns = not significant).

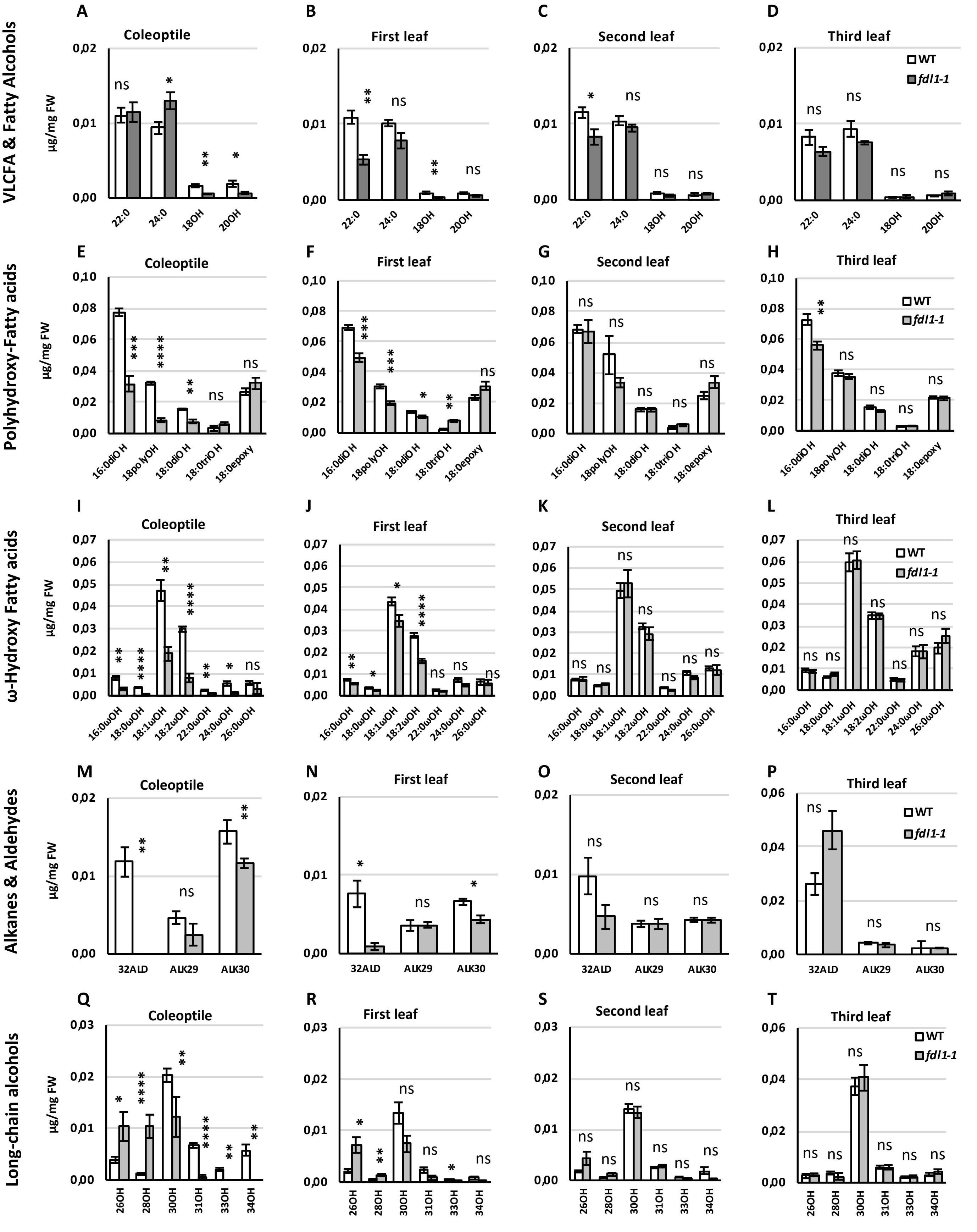


**Supplemental Figure S2. Detailed cuticle composition in seedlings at succeeding developmental stages**. Relative amounts of (A-D) fatty acids (including very long chain fatty acids), (E-H) polyhydroxy-fatty acids, (I-L) ω-hydroxy fatty acids, (M-P) alkanes and aldehydes, and (Q-T) minor long-chain primary alcohols in wild type (WT) and homozygous *fdl1-1* mutant plants. Data are referred to seedlings at coleoptile (A, E, I, M and Q), first leaf (B, F, J, N and R), second leaf (C, G, K, O and S) and third leaf (D, H, L, P and T) developmental stage. Compounds A to L and M to T represent cutin and wax monomers, respectively. Values represent the mean ± SE of five biological replicates. Significant differences were assessed by Student's t-test (* = P < 0.05, ** = P < 0.01, *** = P < 0.001, **** = P < 0.0001, ns = not significant).

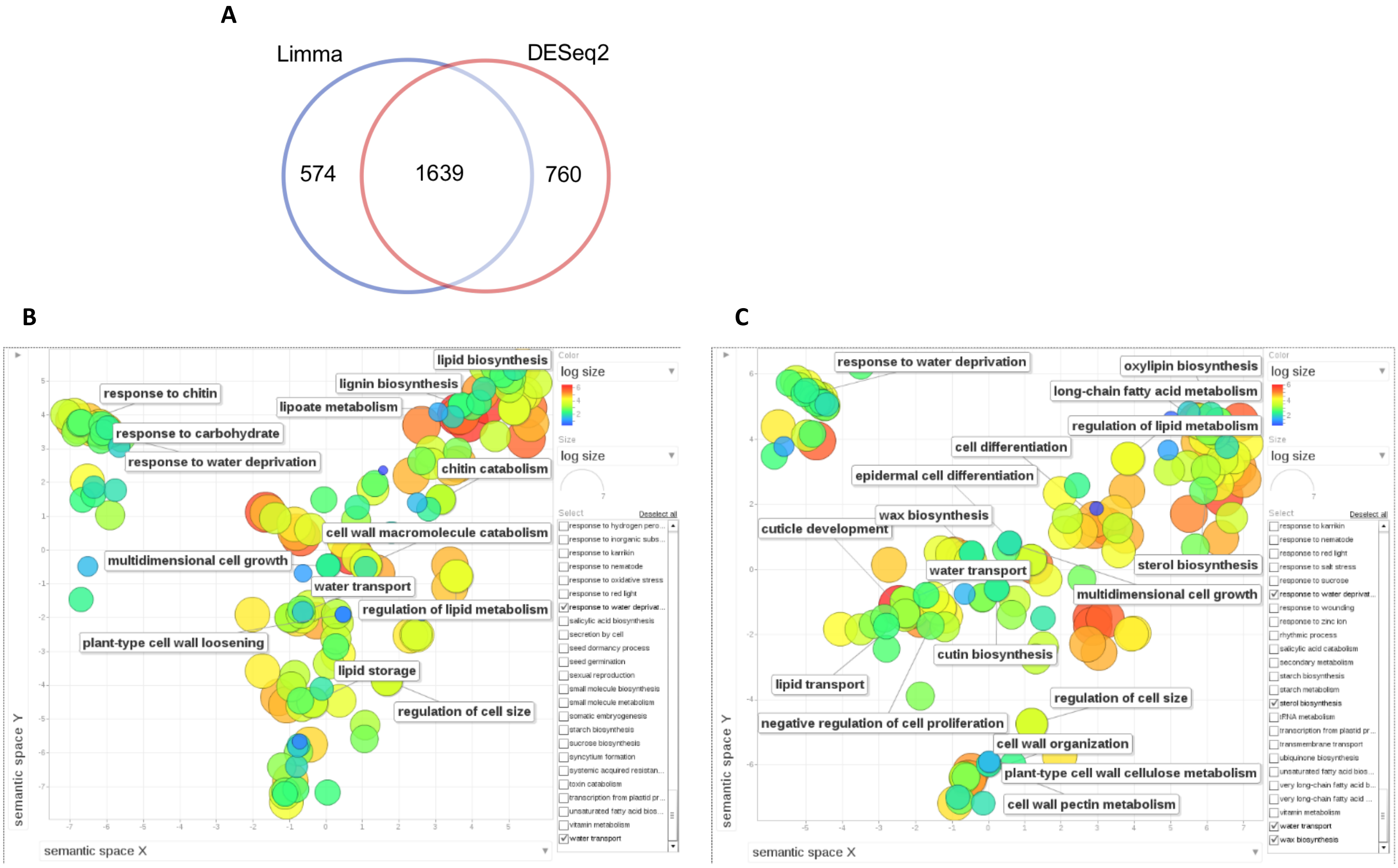


**Supplemental Figure S3. Venn diagram of Differentially expressed genes and representation of enriched gene ontology terms, according to the REViGO tool.** A) Venn diagram of differentially expressed genes, according to Limma (blue) and DESeq2 (red). B) REViGO graphical summary of Gene Ontology terms enriched in the up-regulated genes. Colours are used to indicate the level of significance of the enrichment. Ontology terms relative to wax-biosynthesis, cutin, water transport and lipid metabolism are highlighted. C) REViGO graphical summary of Gene Ontology terms enriched in the down-regulated genes. Colours are used to indicate the level of significance of the enrichment. Ontology terms relative to wax-biosynthesis, cutin, water transport and lipid metabolism are highlighted.

### Supplemental Table S1.

| Pathway | *Zea mays* Gene ID | Gene symbol | Log2 FC | FDR-DESeq2 | FDR-limma | Annotation | *Oryza stiva* Gene ID | *Arabidopsis thaliana* Gene ID | Gene symbol |
|---|---|---|---|---|---|---|---|---|---|
| Lipid metabolism | Zm00001d010451 | *GELP10* | -0.5 | 0.036 | 0.008 | GDSL-type esterase/lipase protein | Os05g0209600 | AT5G45910 | *GELP10* |
| | Zm00001d011269 | *GELP10* | 1.7 | 0.000 | 0.019 | GDSL-type esterase/lipase protein | Os06g0725100 | AT5G55050 | *GELP10* |
| | Zm00001d014918 | *GELP41/EXL3* | -0.9 | 0.000 | 0.009 | GDSL-type esterase/lipase protein | Os02g0101400 | AT1G75900 | *GELP41/EXL3* |
| | Zm00001d016031 | *GELP10* | 0.8 | 0.000 | 0.004 | GDSL-type esterase/lipase protein | Os06g0157000 | AT5G45910 | *GELP10* |
| | Zm00001d017989 | *GELP35* | 1.3 | 0.040 | 0.011 | GDSL-type esterase/lipase protein | Os02g0740400 | AT1G71250 | *GELP35* |
| | Zm00001d019985 | *GELP10* | 1.3 | 0.002 | 0.006 | GDSL-type esterase/lipase protein | Os06g0725100 | AT5G55050 | *GELP10* |
| | Zm00001d031814 | *LCAT/PLA* | -3.9 | 0.000 | 0.015 | lipid acylhydrolase | Os08g0477100 | AT2G26560 | *LCAT/PLA/PLP2* |
| | Zm00001d035865 | acyl-transferases | 1.8 | 0.009 | 0.045 | acyl-CoA dependent acyltransferases | Os06g0103200 | AT5G42830 | nd |
| | Zm00001d036175 | *GELP67* | -1.7 | 0.000 | 0.027 | GDSL-type esterase/lipase protein | Os06g0149100 | AT3G16370 | *GELP67* |
| | Zm00001d036241 | *GELP10* | 1.1 | 0.000 | 0.003 | GDSL-type esterase/lipase protein | Os06g0157000 | AT5G45910 | *GELP10* |
| | Zm00001d038143 | *GELP26* | 0.4 | 0.035 | 0.014 | GDSL-type esterase/lipase protein | Os05g0401000 | AT1G54790 | *GELP26* |
| | Zm00001d039529 | *GELP86* | 1.5 | 0.000 | 0.004 | GDSL-type esterase/lipase protein | Os09g0567800 | AT5G03610 | *GELP86* |
| | Zm00001d039535 | acyl-transferases | 0 | 0.021 | 0.040 | acyl-CoA dependent acyltransferases | Os01g0185300 | AT3G62160 | nd |
| | Zm00001d043680 | *LCAT/PLA* | 0.8 | 0.017 | 0.027 | lipid acylhydrolase | Os01g0710700 | AT2G42690 | *LCAT/PLA* |
| | Zm00001d051419 | *GELP10* | 0.6 | 0.014 | 0.005 | GDSL-type esterase/lipase protein | Os02g0669000 | AT5G55050 | *GELP10* |
| Glycerolipid metabolism | Zm00001d002163 | *DGK5* | -0.8 | 0.002 | 0.013 | diacylglycerol kinase | Os04g0634700 | AT2G20900 | *DGK5* |
| | Zm00001d003750 | *DGD1* | 0.9 | 0.000 | 0.004 | digalactosyldiacylglycerol synthase | Os04g0416900 | AT3G11670 | *DGD1* |
| | Zm00001d008600 | *SQD2* | -1.1 | 0.002 | 0.015 | sulfoquinovosyltransferase | Os01g0142300 | AT5G01220 | *SQD2* |
| | Zm00001d018333 | *DGK7* | -0.4 | 0.013 | 0.030 | diacylglycerol kinase | Os02g0787800 | AT4G30340 | *DGK7* |
| | Zm00001d018406 | *MGD2* | -0.5 | 0.015 | 0.007 | 1,2-diacylglycerol 3-beta-galactosyltransferase | Os02g0802700 | AT5G20410 | *MGD2* |
| | Zm00001d022445 | *AGAL* | -0.6 | 0.019 | 0.014 | alpha-galactosidase | Os07g0679300 | AT3G56310 | *AGAL* |
| | Zm00001d032608 | *AGAL1* | 3.2 | 0.000 | 0.006 | alpha-galactosidase | Os10g0492900 | AT5G08380 | *AGAL1* |
| | Zm00001d033797 | *MAGL4* | 3.3 | 0.000 | 0.009 | acylglycerol lipase | Os03g0719400 | AT1G73480 | *MAGL4* |
| | Zm00001d036982 | *LN1/TAG1* | 0.9 | 0.000 | 0.037 | membrane bound O-acyl transferase | Os06g0563900 | AT2G19450 | *TAG1* |
| Glycerophospholipid metabolism | Zm00001d005383 | *LCAT1* | 2.5 | 0.000 | 0.039 | phospholipid:diacylglycerol acyltransferase | Os02g0590400 | AT1G27480 | *LCAT1* |
| | Zm00001d005392 | *LCAT1* | 0.9 | 0.000 | 0.050 | phospholipid:diacylglycerol acyltransferase | Os02g0590400 | AT1G27480 | *LCAT1* |
| | Zm00001d015399 | *CCT1* | 0.6 | 0.000 | 0.010 | choline-phosphate cytidylyltransferase | Os02g0173500 | AT2G32260 | *CCT1* |
| | Zm00001d019218 | *GPDH5* | -0.8 | 0.001 | 0.019 | glycerol-3-phosphate dehydrogenase | OS07G0229800 | AT2G40690 | *GLY1/SFD1* |
| | Zm00001d040205 | *NPC6* | 0.6 | 0.046 | 0.037 | Phosphoric-diester hydrolases | Os01g0102000 | AT3G48610 | *NPC6* |
| Acetyl-CoA biosynthesis | Zm00001d004125 | *ACC2* | -0.5 | 0.011 | 0.019 | acetyl-CoA carboxylase | Os05g0295300 | AT1G36180 | *ACC2* |
| | Zm00001d009212 | *LPD1* | -0.4 | 0.025 | 0.038 | dihydrolipoamide dehydrogenase | Os01g0337900 | AT3G16950 | *LPD1* |
| | Zm00001d040603 | *ACLB-1* | -0.4 | 0.015 | 0.006 | ATP-citrate lyase | Os01g0300200 | AT3G06650 | *ACLB-1* |
| | Zm00001d043376 | *AAE13* | -0.9 | 0.000 | 0.003 | malonyl-CoA/methylmalonyl-CoA synthetase | Os01g0761300 | AT3G16170 | *AAE13* |
| | Zm00001d048627 | *ACLA-3* | -0.3 | 0.030 | 0.025 | ATP-citrate lyase | Os11g0696200 | AT1G09430 | *ACLA-3* |
| Fatty acid biosynthesis | Zm00001d004019 | *SSI2/FAB2* | 0.3 | 0.046 | 0.006 | acyl-[acyl-carrier-protein] desaturase | Os04g0379900 | AT2G43710 | *SSI2/FAB2* |
| | Zm00001d006866 | *FAB1.1* | -0.4 | 0.025 | 0.016 | 3-oxoacyl-[acyl-carrier-protein] synthase II | Os07g0616200 | AT1G74960 | *KAS2/FAB1* |
| | Zm00001d017418 | *ALDH3F1* | 2.5 | 0.000 | 0.017 | aldehyde dehydrogenase 3F1 | Os02g0647900 | AT4G36250 | *ALDH3F1* |
| | Zm00001d022144 | *FAB1.2* | -0.9 | 0.000 | 0.008 | 3-oxoacyl-[acyl-carrier-protein] synthase II | Os07g0616200 | AT1G74960 | *KAS2/FAB1* |
| | Zm00001d041701 | *ACP4* | -0.4 | 0.015 | 0.012 | acyl carrier protein | Os12g0534200 | AT4G25050 | *ACP4* |
| | Zm00001d047743 | *FAD7* | -1.3 | 0.000 | 0.006 | acyl-lipid omega-3 desaturase | Os03g0290300 | AT5G05580 | *FAD8* |
| Fatty acid elongation | Zm00001d008622 | *GL26* | -0.8 | 0.000 | 0.003 | enoyl-CoA reductase | Os01g0150000 | AT3G55360 | *CER10* |
| | Zm00001d029350 | *KCS24* | -1.5 | 0.003 | 0.012 | 3-ketoacyl-CoA synthase | Os03g0382100 | AT2G28630 | *KCS12* |
| | Zm00001d033637 | *ELO1* | 0.9 | 0.008 | 0.006 | 3-ketoacyl-CoA synthase | OS03G0701500 | AT3G06470 | *ELO2* |
| | Zm00001d037328 | *KCS17* | -1.0 | 0.017 | 0.005 | 3-ketoacyl-CoA synthase | Os06g0260500 | AT1G68530 | *KCS6/CER6* |
| | Zm00001d039053 | *KCS15* | 0.9 | 0.000 | 0.024 | 3-ketoacyl-CoA synthase | Os05g0568000 | AT1G19440 | *KCS4* |
| | Zm00001d039856 | *PAS2* | -0.5 | 0.000 | 0.011 | hydroxyacyl-CoA Dehydratase | Os04g0271200 | AT5G10480 | *PAS2* |
| | Zm00001d044579 | *KCS22/CER60** | -0.8 | 0.011 | 0.054* | 3-ketoacyl-CoA synthase | Os01g0529800 | AT1G25450 | *KCS5/CER60* |
| Cutin and wax biosynthesi | Zm00001d004817 | *FAAH* | -1.2 | 0.000 | 0.006 | fatty acid amide hydrolase | Os11g0170000 | AT5G64440 | *FAAH* |
| | Zm00001d009182 | *ECH1* | -0.3 | 0.009 | 0.020 | 3-hydroxyacyl-CoA dehydrogenase | Os01g0348600 | AT3G06860 | *MFP2* |
| | Zm00001d010855 | *LDAP3* | 0.6 | 0.000 | 0.040 | rubber elongation factor protein | Os05g0151300 | AT3G05500 | *LDAP3* |
| | Zm00001d017251 | *CER1* | 1.0 | 0.000 | 0.004 | aldehyde decarbonylase | Os02g0621300 | AT1G02205 | *CER1* |
| | Zm00001d020238 | *ONI3* | -0.9 | 0.000 | 0.025 | fatty acid omega-hydroxy dehydrogenase | Os09g0363900 | AT1G72970 | *EDA17* |
| | Zm00001d031893 | *HCT12/DCR* | -0.8 | 0.000 | 0.008 | diacylglycerol acyltransferase | OS08G0562500 | AT5G23940 | *PEL3/DCR* |
| | Zm00001d032284 | *HTH1* | -0.8 | 0.004 | 0.006 | fatty acid omega-hydroxy dehydrogenase | Os08g0401500 | AT1G72970 | *EDA17* |
| | Zm00001d034319 | *FAH1* | -0.5 | 0.037 | 0.018 | fatty acid 2-hydroxylase | Os03g0780800 | AT2G34770 | *FAH1* |
| | Zm00001d034832 | *AAE16* | -0.5 | 0.000 | 0.006 | long-chain-alcohol oxidase | Os03g0845500 | AT3G23790 | *AAE16* |
| | Zm00001d044136 | *GPAT8* | 1.2 | 0.036 | 0.032 | acyl-CoA:sn-glycerol-3-phosphate 1-O-acyltransferase | Os01g0631400 | AT1G06520 | *GPAT1* |
| | Zm00001d045295 | *LACS4* | -0.7 | 0.000 | 0.032 | long-chain acyl-CoA synthetase | Os06g0158000 | AT4G23850 | *LACS4* |
| | Zm00001d049950 | *MSH1/CER4* | -1.1 | 0.000 | 0.003 | alcohol-forming fatty acyl-CoA reductase | Os08g0557800 | AT4G33790 | *FAR3/CER4* |
| | Zm00001d051130 | *FAO4* | -0.7 | 0.044 | 0.013 | long-chain-alcohol oxidase | Os02g0621800 | AT4G28570 | *FAO4b* |
| | Zm00001d051923 | *CUS2* | -1.1 | 0.009 | 0.006 | cutin synthase 2 | Os02g0816200 | AT5G33370 | *CUS2* |
| | Zm00001d053127 | *LACS2* | -0.9 | 0.000 | 0.009 | long-chain acyl-CoA synthetase | Os11g0558300 | AT1G49430 | *LACS2* |
| | Zm00001d024723 | *CER8** | -1.2 | 0.082* | 0.000 | acyl-CoA synthetase | OS05G0132100 | AT2G47240 | *LACS1/CER8* |
| | Zm00001d043853 | *WSD11** | 3.1 | 0.085* | 0.007 | wax ester synthase | Os01g0681000 | AT5G53390 | *FOP1/WSD11* |
| Fatty acid, cutin and wax transporter | Zm00001d005146 | *DIR1* | 1.8 | 0.000 | 0.011 | apoplastic lipid transfer protein | Os07g0287400 | AT5G48485 | *LTP/DIR1* |
| | Zm00001d010426 | *CER5* | -0.8 | 0.001 | 0.040 | ABC transporter | Os05g0222200 | AT1G51500 | *ABCG12/CER5* |
| | Zm00001d011091 | *CTS* | -0.6 | 0.000 | 0.005 | peroxisomal ABC transporter 1 | Os01g0966100 | AT4G39850 | *ABCD1/CTS* |
| | Zm00001d012535 | *LTPG2* | -1.1 | 0.000 | 0.003 | lipid transport | Os05g0489200 | AT3G43720 | *LTPG2* |
| | Zm00001d013960 | *ABCG11* | -0.6 | 0.015 | 0.014 | ABC transporter | Os10g0494300 | AT1G17840 | *ABCG11/COF1* |
| | Zm00001d018278 | *CHI4/FAP2* | 0.4 | 0.025 | 0.029 | fatty acid binding protein | Os02g0778500 | AT2G26310 | *FAP2* |
| | Zm00001d034513 | *LTPG6* | -1.4 | 0.000 | 0.003 | lipid transport | Os03g0804200 | AT1G55260 | *LTPG6* |
| | Zm00001d039631 | *GL13* | -0.9 | 0.000 | 0.010 | ABC transporter | Os02g0208300 | AT2G26910 | *PDR4* |
| | Zm00001d043049 | *PLT3* | -0.4 | 0.029 | 0.008 | lipid transport | Os01g0822900 | AT5G01870 | *LTP* |
| | Zm00001d048718 | *ACBP4* | -0.7 | 0.001 | 0.011 | acyl-CoA binding protein | Os11g0657000 | AT3G05420 | *ACBP4* |
| Regulatory proteins | Zm00001d020457 | *MYB30* | -1.1 | 0.000 | 0.004 | MYB transcription factor | Os08g0437300 | AT3G28910 | *MYB30* |
| | Zm00001d026486 | *SHN2.2* | -1.5 | 0.004 | 0.024 | SHINE Clade/AP2 Domain Transcription Factors | Os04g0655700 | AT5G25190 | *SHN2* |
| | Zm00001d035835 | *SHN2.1* | 2.7 | 0.000 | 0.007 | SHINE Clade/AP2 Domain Transcription Factors | Os06g0181700 | AT5G25190 | *SHN2* |
| | Zm00001d040090 | *OCL1* | -0.6 | 0.000 | 0.012 | MYB transcription factor | Os02g0674800 | AT4G00730 | *ANL2* |
| | Zm00001d046621 | *GL15* | -1.3 | 0.000 | 0.014 | AP2 transcription factor | Os03g0818800 | AT4G36920 | *FLO2* |

**Supplemental Table S1. Differentially expressed genes involved in cuticle-related processes in the *fdl1-1* mutant.** List of *Zea mays* gene IDs putatively involved in lipid, cutin and wax metabolism. Gene symbols, fold change (FC), corrected P values (FDR) and annotations are reported. Asterisks indicate genes with significative P value only for one of the two method used to find DEGs. Gene IDs of the closest orthologues in *Oryza sativa* and *Arabidopsis thaliana* are also listed.

| *Zea mays* | | | | | | *Oryza sativa* | *Arabidopsis thaliana* | |
|---|---|---|---|---|---|---|---|---|
| **Gene ID** | **Gene symbol** | **Log2 FC** | **FDR-DESeq2** | **FDR-limma** | **Annotation** | **Gene ID** | **Gene ID** | **Gene symbol** |
| Zm00001d002288 | *BHLH148* | 1.1 | 9.4E-04 | 9.3E-03 | bHLH-transcription factor | Os04g0618600 | AT2G20180 | *PIL5/PIF1* |
| Zm00001d051569 | *BHLH11* | 0.7 | 2.3E-02 | 1.6E-02 | bHLH-transcription factor | Os02g0705500 | AT4G34530 | *CIB1* |
| Zm00001d018056 | *BHLH30* | -0.8 | 7.9E-07 | 6.0E-03 | bHLH-transcription factor | Os02g0747900 | AT1G26945 | *KDR* |
| Zm00001d044599 | *PHOT1* | -1.3 | 4.3E-09 | 6.8E-03 | blue light receptor | Os11g0102200 | AT3G45780 | *PHOT1* |
| Zm00001d032353 | *PHOT2* | 1.4 | 1.7E-03 | 3.7E-03 | blue light receptor | Os04g0304200 | AT5G58140 | *PHOT2* |
| Zm00001d008734 | *bZIP122* | -1.5 | 5.4E-13 | 6.1E-03 | bZIP-transcription factor | Os01g0174000 | AT5G11260 | *HY5* |
| Zm00001d015743 | *bZIP61* | -1.3 | 1.3E-08 | 3.2E-02 | bZIP-transcription factor | Os02g0202950 | AT5G11260 | *HY5* |
| Zm00001d039658 | *bZIP6* | -0.9 | 1.4E-04 | 6.3E-03 | bZIP-transcription factor | Os01g0174000 | AT5G11260 | *HY5* |
| Zm00001d045944 | *CRY3* | -0.4 | 7.9E-03 | 4.6E-02 | cryptochrome | Os02g0625000 | AT1G04400 | *CRY2* |
| Zm00001d007445 | *FKF1* | -1.5 | 2.1E-05 | 7.4E-03 | flavin-binding kelch repeat f-box | Os11g0547000 | AT1G68050 | *ADO3/FKF1* |
| Zm00001d053091 | *FKF2* | -1.9 | 5.7E-06 | 4.7E-03 | flavin-binding kelch repeat f-box | Os11g0547000 | AT1G68050 | *ADO3/FKF1* |
| Zm00001d008826 | *GI1* | -0.9 | 1.3E-08 | 9.0E-03 | gigantea protein | Os01g0182600 | AT1G22770 | *GI* |
| Zm00001d039589 | *GI2* | -1.4 | 6.7E-24 | 3.1E-03 | gigantea protein | Os01g0182600 | AT1G22770 | *GI* |
| Zm00001d024546 | *LHY1* | 1.1 | 6.9E-05 | 2.7E-02 | MYB transcription factor | Os08g0157600 | AT1G01060 | *LHY1* |
| Zm00001d049543 | *CCA1* | 3.0 | 6.8E-24 | 6.5E-03 | MYB transcription factor | Os08g0157600 | AT1G01060 | *LHY1* |
| Zm00001d013402 | *PHYA2* | -0.9 | 6.7E-05 | 5.9E-03 | red/far-red light receptor | Os03g0719800 | AT1G09570 | *PHYA* |
| Zm00001d043592 | nd | -1.0 | 6.5E-07 | 3.3E-02 | suppressor of phythchrome A | Os01g0725800 | AT3G15354 | *SPA3* |
| Zm00001d039072 | nd | -0.4 | 2.3E-02 | 1.5E-02 | suppressor of phythchrome A | Os05g0571000 | AT4G11110 | *SPA2* |
| Zm00001d047761 | nd | -0.5 | 9.8E-04 | 7.5E-03 | two-component response regulator | Os03g0284100 | AT5G60100 | *PRR3* |
| Zm00001d004875 | *PRR59* | -0.9 | 2.2E-08 | 7.3E-03 | two-component response regulator | Os11g0157600 | AT2G46670 | *PRR5* |
| Zm00001d006212 | *PRRH1* | -3.3 | 8.0E-44 | 3.7E-03 | two-component response regulator | Os09g0532400 | AT2G46790 | *PRR9* |
| Zm00001d052781 | nd | -1.5 | 9.0E-18 | 1.2E-02 | two-component response regulator | Os11g0157600 | AT2G46790 | *PRR9* |
| Zm00001d022590 | *PRR37a* | -0.6 | 7.3E-03 | 1.2E-02 | two-component response regulator | Os07g0695100 | AT5G02810 | *PRR7* |
| Zm00001d021291 | nd | -3.1 | 7.1E-66 | 2.1E-03 | two-component response regulator | Os09g0532400 | AT5G24470 | *PRR5* |
| Zm00001d017241 | *TOC2* | -1.9 | 4.4E-17 | 1.5E-02 | two-component response regulator | Os02g0618200 | AT5G61380 | *TOC1* |
| Zm00001d051114 | *TOC1* | -0.9 | 1.7E-05 | 8.3E-03 | two-component response regulator | Os02g0618200 | AT5G61380 | *TOC1* |
| Zm00001d039156 | nd | -0.6 | 5.9E-03 | 6.1E-03 | unknown | Os01g0566100 | AT2G25930 | *ELF3* |
| Zm00001d046935 | nd | -1.1 | 5.7E-07 | 1.2E-02 | unknown | Os06g0661800 | AT5G24850 | *CRY3* |
| Zm00001d042935 | nd | -1.1 | 9.3E-03 | 2.2E-02 | unknown | Os06g0661800 | AT5G24850 | *CRY3* |

**Supplemental Table S2. Differentially expressed genes involved in circadian rhythm in the *fdl1-1* mutant.** List of *Zea mays* gene IDs involved in photoreception and circadian rhythm pathway. Gene symbols, fold change (FC), corrected P values (FDR) and annotations are reported. Gene IDs of the closest orthologues in *Oryza sativa* and *Arabidopsis thaliana* are also listed.

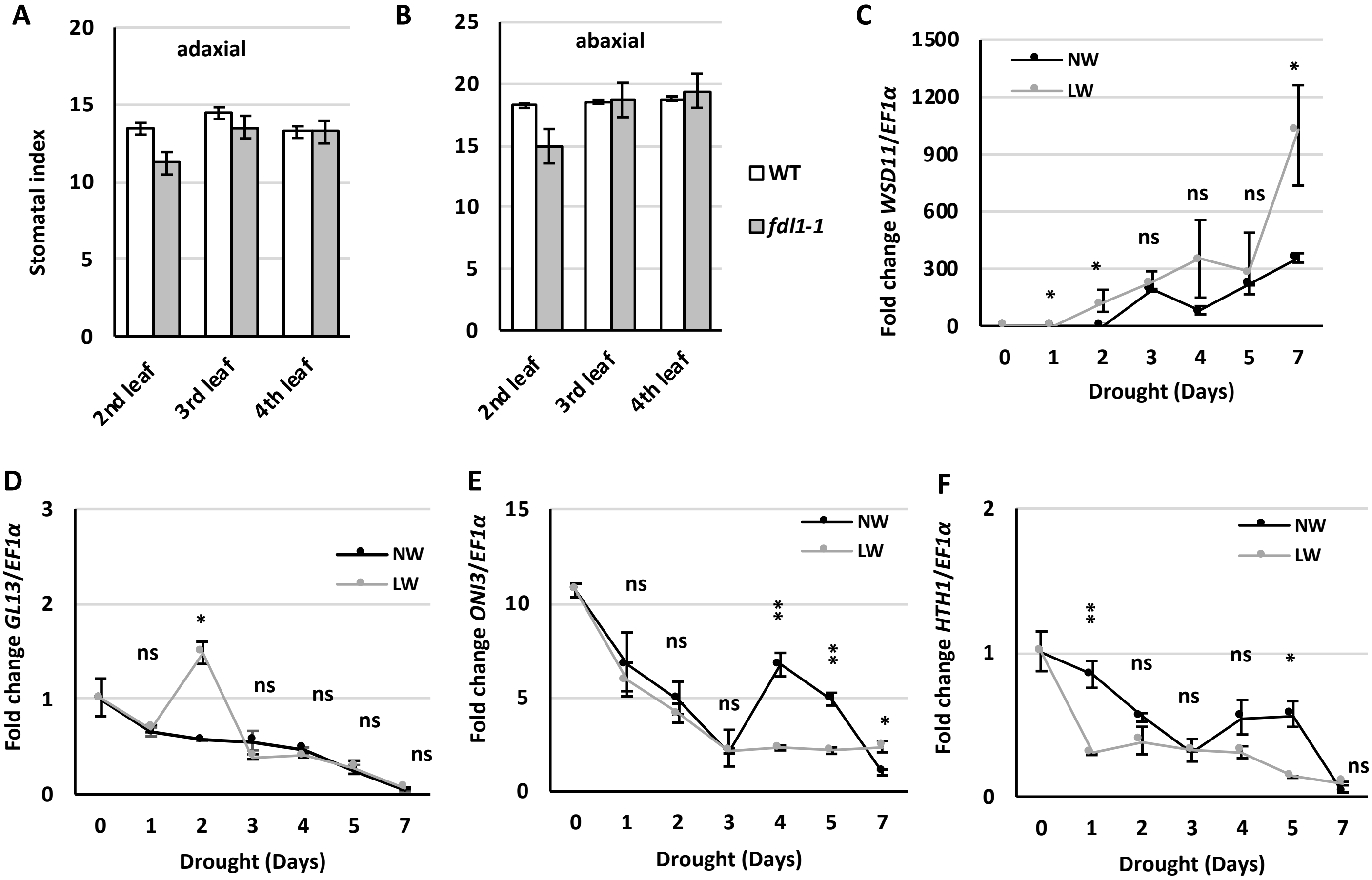


**Supplemental Figure S4. Stomatal index and additional information about drought stress experiments.** Stomatal index measured on the adaxial (A) and the abaxial (B) side of the second, third and fourth leaf of *fdl1-1* homozygous and wild type (WT) plants. Values are the mean ± SE of ten replicates. Comparison has been made between wild type and *fdl1-1* genotypes. (C-F) Transcript accumulation of the putative cuticle related *ZmWSD11, ZmGL13*, *ZmONI3* and *ZmHTH1* genes, analysed for 7 days by qRT-PCR, in wild type second leaf of plants grown under low water (LW) or normal (NW) conditions. Values represent the mean fold change variations ± SD of three biological replicates. Comparison was made at each time point between LW and control (NW) plants. Significant differences were assessed by Student's t-test (* = $P < 0.05$, ** = $P < 0.01$, *** = $P < 0.001$, **** = $P < 0.0001$, ns = not significant).

**Supplemental Table S3. Primers used for Real Time PCR analysis.**

| Gene ID | Gene Name | Forward Sequence (5'- 3') | Reverse Sequence (5'- 3') | References |
|---|---|---|---|---|
| Zm00001d046449 | EF1α | TGGGCCTACTGGTCTTACTACTG | ACATACCCACGCTTCAGATCCT | Lin et al. 2014 |
| Zm00001d045321 | RAB18 | TATGGGACGACGACCACTGAA | TAGTGCTGACCGGGGAGTTT | |
| Zm00001d022227 | FDL1 | TAGCTGTTCAGATCGGTCG | CCACACAACATGCAACTTGC | La Rocca et al. 2016 |
| Zm00001d049950 | CER4 | CGGCTCTACAACGACCTCAA | CCTTAGCCGCTCCAGGTTTA | |
| Zm00001d017251 | CER1 | TGTGGTGGTACATGTGGGTG | CAGGCCGTACTGGAAGTTGT | |
| Zm00001d039631 | GL13 | ACCATTGCGCCTATTATTGC | CCGAACTGAGAGGTCAGGAG | Li et al. 2013 |
| Zm00001d046621 | GL15 | GGTAAAAGCAGTGGCAGAGG | TAACTAGTGGCCACCCCAAG | |
| Zm00001d032284 | HTH1 | TACCCAAGCACACAGACGAC | CCCCCATGTATCTGCCCATC | |
| Zm00001d020238 | ONI3 | CTGCTGATGGTGCTGGGTTA | GGTTGGTCCCTGGCGATTTA | |
| Zm00001d043853 | WSD11 | TCTAGCGCTCTGTCTCGGTA | AGCATGTAGCCCAGCTTGTT | |
| Zm00001d003190 | | GCGAACGTGGCTAATGTGG | TCCGGGTGTAGAAGTTCTTGC | |
| Zm00001d030316 | | TTCAAGGGTCTTGAGGTGGC | CGTGCTAAGTGGGATGCTGA | |
| Zm00001d048947 | | CGTACCACCTGTACAACCCG | CGCACGGTTCTTCACCCTTA | |
| Zm00001d031158 | | TCTACGACATCTCGGTCATCG | CACATGCATGGGTCTTCATGTC | |
| Zm00001d038891 | | TTTGTGGGCATGCTCGATCT | CAGAGCACTTGCGTCCAATG | |
